# The Split-Belt Rimless Wheel

**DOI:** 10.1101/2021.04.30.442053

**Authors:** Julia K. Butterfield, Surabhi N. Simha, J. Maxwell Donelan, Steven H. Collins

**Affiliations:** Department of Mechanical Engineering, Stanford University, Stanford, CA, USA; Department of Biomedical Physiology and Kinesiology, Simon Fraser University, Burnaby, British Columbia, Canada

**Keywords:** Split-belt treadmill, rimless wheel, energy harnessing, legged locomotion, human walking

## Abstract

Split-belt treadmill training, common in stroke rehabilitation and motor learning experiments, reveals a mechanism through which energy can be extracted from the environment. People can extract net positive work from a split-belt treadmill by lengthening their step onto the fast belt. To understand how leg angles and belt speed differences affect energy transfer between the treadmill and the person during split-belt walking, we simulated a split-belt rimless wheel that alternates rotating on fast and slow treadmill belts. We found that the split-belt rimless wheel can passively walk steadily forward under a range of conditions, extracting enough energy from the treadmill to overcome losses during collisions. The simulated wheel can tolerate both speed disturbances and ground height variability, and it can even capture enough energy to walk uphill. We also built a physical split-belt rimless wheel robot, demonstrating the feasibility of energy extraction during split-belt treadmill walking. In comparing the wheel solutions to human split-belt gait, we found that humans do not maximize positive work performed by the treadmill; costs associated with balance and free vertical moments likely limit adaptation. This study characterizes the mechanics and energetics of split-belt walking, demonstrating that energy capture through intermittent contacts with the two belts is possible when the belt speed difference is paired with an asymmetry in leg angles at step-to-step transitions. This study demonstrates a novel way of harnessing energy through individual rotations rather than continuous contact and offers a simple model framework for understanding human choices during split-belt walking.

## 1 Introduction

Split-belt treadmills are commonly used for stroke rehabilitation and motor learning experiments (Reisman et al.2010; Roemmich and Bastian 2018). Individuals post-stroke often adopt an asymmetric gait (Finley et al. 2015; Patterson et al. 2008; Wall and Turnbull 1986), and split-belt treadmill training has been proposed and applied to help individuals relearn a symmetric gait (Helm and Reisman 2015; Reisman et al. 2010; Roemmich and Bastian 2018). Although many effects of repeated split-belt training remain unexplored, this training has moved beyond clinical trials and is currently being implemented in some physical therapy clinics (Beyaert et al. 2015; Patterson et al. 2015; Marianjoy Rehabilitation Hospital 2016). Researchers are even working to make splitbelt training portable so that people who have had a stroke can train outside the confines of the lab (Aucie et al. 2020; Kim et al. 2019).

Researchers also use the split-belt treadmill paradigm to understand how people adapt to novel motor environments (Reisman et al. 2010; Roemmich and Bastian 2018) and how adaptation can differ between healthy individuals and those with gait pathologies such as amputation (Kline et al. 2020; Selgrade et al. 2017b), cerebral palsy (Levin et al. 2017; Mawase et al. 2016), and Parkinson’s disease (Seuthe et al.2019). The field of split-belt treadmill research continues to grow as researchers investigate spatiotemporal adaptation (Finley et al. 2015; Stenum and Choi 2020; Yokoyama et al.2018) and explore how energy minimization contributes to observed changes in gait (Finley et al. 2013; Sánchez et al.2017, 2019, 2020; Stenum and Choi 2020). As split-belt treadmills become more prevalent, both for rehabilitation interventions and motor learning experiments, there is a need to learn more about the underlying mechanics of locomotion on split-velocity surfaces.

Recent experimental results have revealed that people can take advantage of positive work performed by a splitbelt treadmill. As people adapt to the treadmill, they tend to lengthen their step onto the fast belt. This longer step allows the treadmill to do positive work on the person, which often correlates with a decreased metabolic energy cost (Sánchez et al. 2019; Selgrade et al. 2017a). Understanding the mechanism through which people capture energy from the treadmill would be useful for understanding what drives split-belt adaptation.

The process through which people extract energy from the split-belt treadmill during walking is different from traditional, industrial methods of energy extraction. Humans have captured energy from velocity differences in the environment for millennia (Temple and Needham 1986), but traditional applications rely on continuous contact with two surfaces of different velocity. For example, water wheels rest on a stationary riverbank while a moving river turns the wheel to generate power (Reynolds 1983). Sailboats exploit the speed difference between the wind and water to propel themselves through the water (Carter 2002). Land yachts use propellers to drive downwind faster than the wind (Bauer 1969; Khan et al. 2013), and a cart with geared wheels can move forward on a split-belt treadmill with both belts moving backward at different speeds (Chiu et al. 2019). People walking on a split-belt treadmill do not continuously contact both belts, however, but instead alternate between the fast and slow belts.

Nature does provide an example of an energy harnessing mechanism that does not rely on continuous contact: the albatross flying with dynamic soaring. Flying in shallow arcs with and against the wind across the boundary layer above the ocean (Bousquet et al. 2017), an albatross can extract energy from the wind to balance its drag losses, enabling steady flight without flapping. The wind speeds are low close to the ocean’s surface and increase as the height above the water increases. As an albatross climbs, it faces slightly into the wind, using the increasing wind speed to gain height. The bird then turns to fly more with the wind as it descends, trading height for speed, before turning to again face into the wind for the next climb (Walkden 1925). While the albatross moves smoothly through a range of wind speeds and humans alternate contacts with the fast and slow treadmill belts, there could be similarities in the mechanisms of energy extraction in these two systems.

Simulations and simple models can be extremely effective in helping us understand complex activities such as human walking. For example, previous walking models helped researchers establish a framework to consider the gravity-driven passive aspects of leg swing (Mochon and McMahon 1980), the types of losses present during walking (Kuo et al.2005), and the possible powering strategies to overcome these losses (Mcgeer 1990). The frameworks from these original models enabled later experimental investigations of the relative contributions of step-to-step transition losses (Donelan et al. 2002) and swing costs (Doke et al. 2005), and analysis of how ankle and hip power contribute to propulsion (Kuo 2002; Lewis and Ferris 2008). Simulation models allow exploration of an entire parameter space in a way not possible in human experiments because of time constraints. Additionally, certain outcomes can be modeled which are difficult or even impossible to measure in experiments.

One common walking model is the rimless wheel, in which a wheel walks forward by rotating on a single spoke and then transitioning onto its next spoke and rotating again. The rimless wheel is simple enough to have analytical solutions, but complete enough to contain both the step-to-step transition and single support phases of walking (Coleman 2010). In normal gait, it captures the kinetic and potential energy exchange of the center of mass motion and reveals the cost of redirecting the center of mass during step-to-step transitions (Coleman et al. 1997). The rimless wheel can be modified for use on the split-belt treadmill, enabling analytical modeling of split-belt walking.

A simple model of split-belt walking must incorporate belt speed differences and adjustable step length asymmetry. Step length asymmetry, known to be associated with changes in energy cost of human gait (Finley et al. 2013), can be achieved in a rimless wheel model with two individual sets of spokes connected with an angular offset. The model’s mass, rotational inertia, and spoke length, as well as the treadmill incline, also have the potential to impact the wheel’s behavior. A modified rimless wheel enables assessment of speed, energy use, and stability during split-belt walking.

In this study, we used a split-belt rimless wheel to model walking on a split-belt treadmill in order to characterize the fundamental features of the mechanics, energetics, and stability of locomotion on a dual-velocity surface. Steady walking for the split-belt rimless wheel is not possible if the belts are moving at the same speed. We hypothesized, however, that under certain split-belt conditions, a split-belt rimless wheel would be able to passively walk forward on a level split-belt treadmill with both belts moving backward by capturing energy from the treadmill to overcome collision losses. We systematically varied the belt speed difference and the angles between the spokes on the split-belt rimless wheel to characterize how asymmetries enable energy capture during split-belt walking. We investigated the impact of the collision angles, belt speed difference, and rotational inertia on the wheel’s average velocity and stability to learn more about why people adopt certain gaits. We simulated uphill slopes, anticipating that the wheel might be able to harness even more energy from the treadmill than is required to walk steadily on level ground. We compared the model solutions to human data for a specific pair of belt speeds to understand which unmodeled costs might limit people’s ability to benefit from a belt speed difference. Finally, we built a working physical prototype to validate the model and demonstrate that a real system can extract energy through short, alternating contacts with different surfaces. We expected the results of this study to guide us toward a better understanding of the ways in which people can extract energy from a splitbelt treadmill, giving us a framework through which to interpret past and future experimental results. Knowledge of the novel mechanism through which intermittent contacts enables energy extraction also creates opportunities for further exploration of energy-capturing devices.

This paper is organized into three main sections. In the first section, we present the split-belt rimless wheel simulation model and describe its dynamics. We introduce the system of equations which must be solvable for the wheel to walk steadily and explain the outcome measures of slow-belt walking speed and disturbance rejection. In the second section, we perform a series of simulations in which we systematically investigate the impacts of the belt speed difference, rotational inertia, treadmill incline, and angles between the spokes at collision onto the fast and slow belts. In the third section, we describe the design of a physical prototype and compare experimental and simulation results of measuring the wheel’s walking speed for a variety of belt speed differences.

## 2 Model Mechanics and Dynamics

### 2.1 Model Definitions and Parameters

The split-belt rimless wheel (Fig. 1) consists of two identical rimless wheels that are rigidly attached. The two sets of spokes are connected with a lateral offset, so that each set only interacts with one belt, and with an angular offset, so that only one spoke touches the ground at any time. The split-belt rimless wheel walks forward by pivoting over one spoke until the next spoke collides with the ground and then pivoting over the new spoke, alternatingly contacting the fast and slow belts. One gait cycle consists of the rotation on the slow belt, the collision onto the fast belt, the rotation on the fast belt, and the collision onto the slow belt. The simulated wheel is constrained to the sagittal plane, with no degrees of freedom for roll or yaw. As a result, the magnitude of the lateral offset between the sets of spokes has no impact on the wheel’s motion.

**Figure 1.**
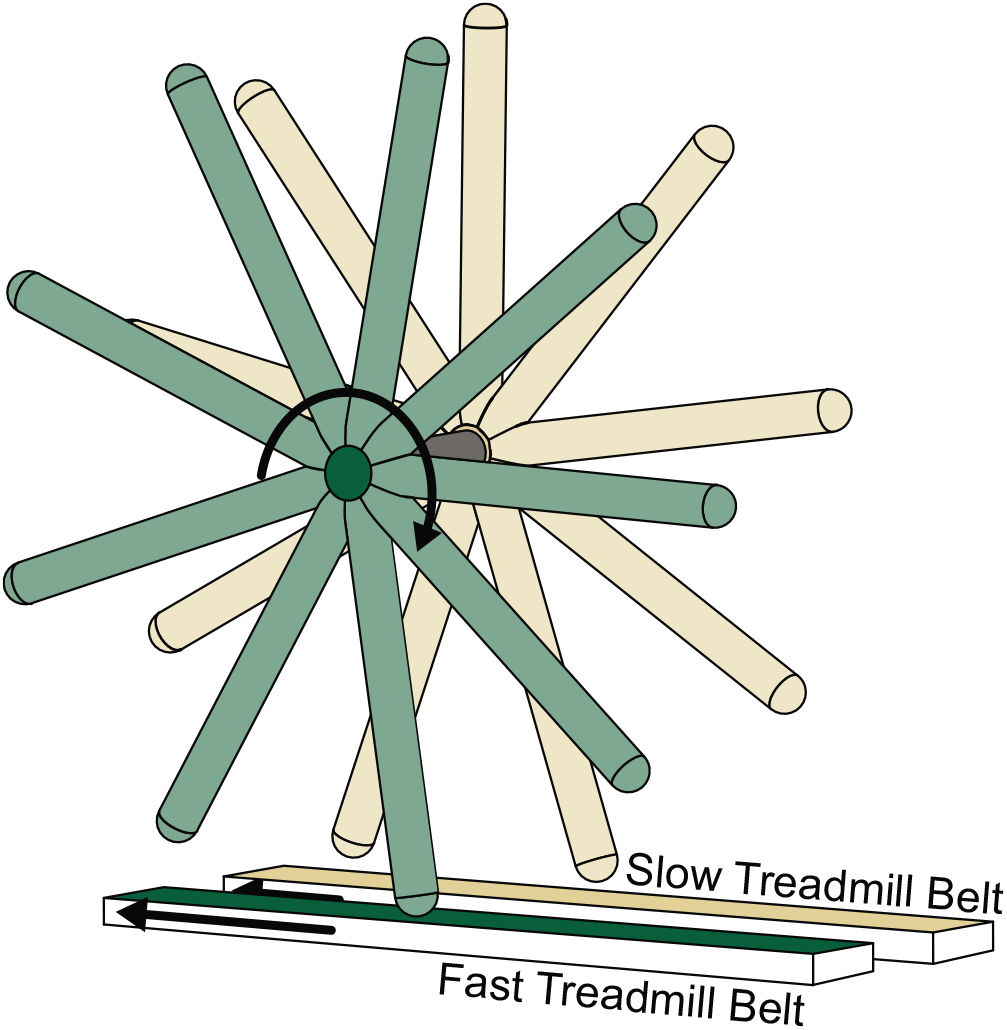
The split-belt rimless wheel consists of two identical rimless wheels that are rigidly attached. The wheel walks forward by pivoting over one spoke until the next spoke collides with the ground and then pivoting over the new spoke. It alternates contacting the fast and slow treadmill belts.

Five parameters define the split-belt rimless wheel: leg length, mass, moment of inertia, and two collision angles (Fig. 2). The leg length *l* is the distance from the center of mass to the end of the spokes. The moment of inertia about the center of mass is 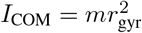 where *m* is the total model mass and *r*_gyr_ is the radius of gyration ranging from 0 with all mass concentrated at the center of the wheel to l with all mass concentrated at the ends of the spokes. The collision angles are the angles between the spokes during the fast and slow belt collisions, called 2*α* and 2*β* respectively. The number of spokes on each wheel and the angular offset between the two sets of spokes together define these collision angles. For a physical model, the sum of 2*α* and 2*β* would need to divide evenly into 360°, but the collision angles can be set arbitrarily in simulation.

**Figure 2.**
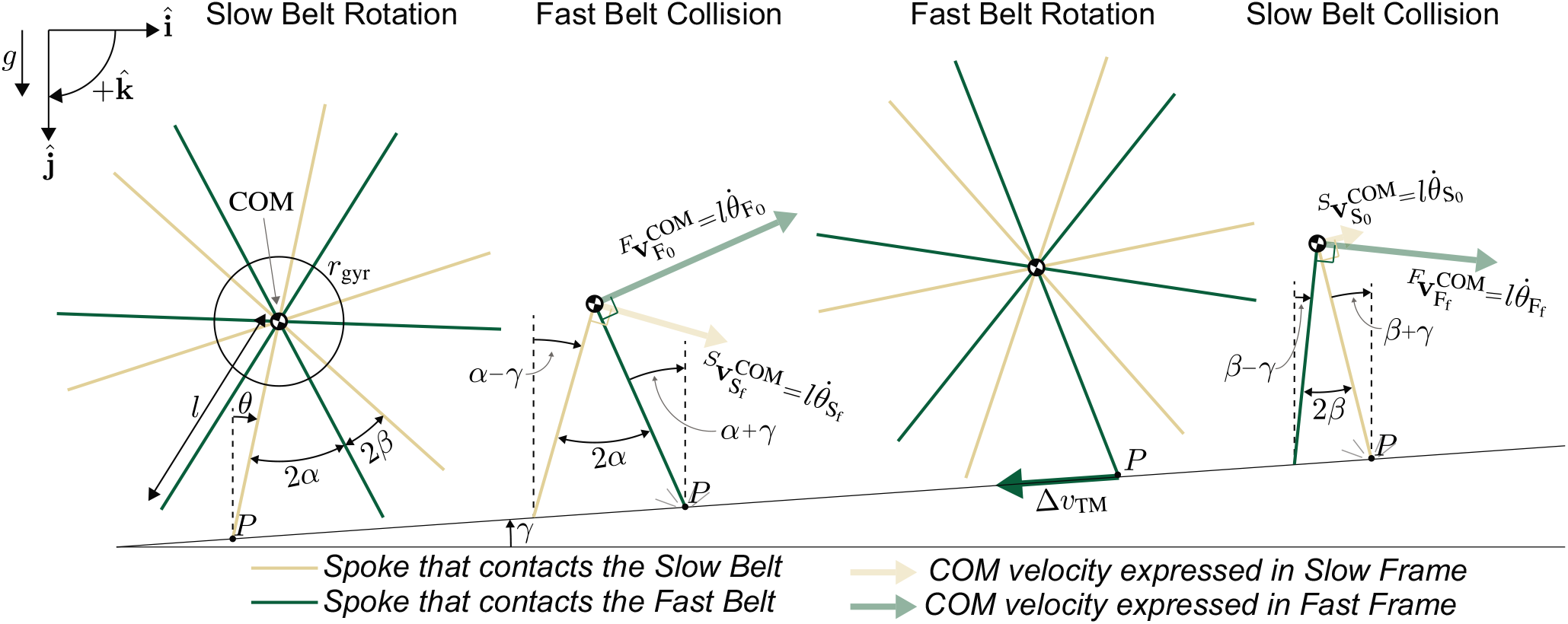
Two-dimensional representation of the split-belt rimless wheel. Collision angles 2*α* and 2*β*, spoke length *l*, and radius of gyration *r*_gyr_ define wheel geometry. The treadmill parameters are the belt speed difference Δν_TM_ and incline *γ*. *P* is the contact point between the wheel and the treadmill, redefined at each collision. Progress through rotations is tracked with *θ*. The velocity of the center of mass **v**^COM^ is always perpendicular to the contacting spoke when expressed in the frame of the contacted belt. The subscripts **S**_0_, **S**_f_, **F**_0_, and **F**_f_ refer to the values at the beginning and end of the slow and fast belt rotations respectively. The left-superscripts *S* and *F* indicate whether the center of mass velocity is shown in the slow or fast belt reference frame.

Three parameters define the split-belt treadmill: belt speed difference, slow belt speed, and treadmill incline. The belt speed difference Δ*ν*_TM_ is the absolute difference between the fast and slow belt speeds. Split-belt treadmill experiments commonly emphasize the ratio between the belt speeds, but the difference, not the ratio, governs performance. The incline *γ* is the angle between the treadmill and level ground.

### 2.2 Reference Frames

There are three distinct inertial reference frames in the system: the stationary lab frame, the slow belt frame, and the fast belt frame. Energies and linear velocities are reference frame-dependent, while forces and angular velocities are frame-independent. Potential energy is only frame-dependent when there is an incline and the wheel moves down while rotating on the moving belts. We annotate frame-dependent quantities with a left-superscript *F* or *S* to indicate whether they are expressed in the fast or slow belt reference frame. We note explicitly any time that we use the stationary lab frame.

Computing or comparing frame-dependent quantities requires expressing the terms in the same reference frame. Linear velocity is transferred from the fast belt frame to the slow belt frame by subtracting out the vector belt speed difference, and from the slow belt frame to the stationary lab frame by subtracting out the slow belt velocity. We use the slow belt reference frame whenever possible. For example, we compute energy in the slow belt frame when tracking wheel energy over the course of a gait cycle. For some calculations, however, such as the energy balance for the fast belt rotation, the fast belt frame is more convenient.

### 2.3 Steady Walking Conditions

We hypothesized that the split-belt rimless wheel could walk steadily on level ground and uphill, with steady walking being defined by the wheel having the same angular velocity at the beginning of consecutive gait cycles. We chose the beginning of the slow belt rotation as the beginning of the cycle, meaning the steady walking condition is

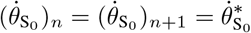

where 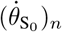 is the angular velocity at the beginning of the *n*th slow belt rotation and 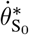 is the fixed point angular velocity.

One equation governs each of the four cycle components, relating the angular velocity before and after each rotation or collision. We label the four key angular velocities 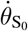, 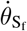, 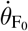, and 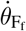 where the subscript S refers to the slow belt rotation and the subscript F refers to the fast belt rotation. The sub-subscript 0 refers to the beginning of the rotation and the sub-subscript f refers to the end of the rotation. The following sections provide a high-level derivation of the equations relating each initial and final angular velocity (a complete derivation can be found in Appendix B).

#### 2.3.1 Rotation on the Slow Belt

Given the velocity at the beginning of the slow belt rotation, the velocity at the end can be determined using an energy balance approach. During the rotation, the split-belt rimless wheel behaves like an inverted pendulum rotating over the spoke in contact with the slow treadmill belt. In the slow belt reference frame, this results in an even exchange of potential and kinetic energy and constant total energy. When solved for the angular velocity at the end of the slow belt rotation 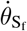, the energy balance is

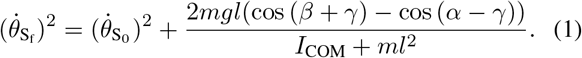

The terms *l* cos (*β* + *γ*) and *l* cos (*α* − *γ*) are the heights of the center of mass at the beginning and end of the rotation respectively, measured relative to the contact point *P* between the spoke and the slow treadmill.

#### 2.3.2 Collision onto the Fast Belt

Conservation of angular momentum about the new foot contact can be used to determine the angular velocity after the fast belt collision 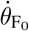. We approximated the collision as instantaneous and impulsive.

The angular momentum **H** in general terms is

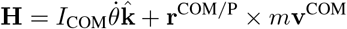

where **r**^COM/P^ is the position vector from the point colliding with the fast treadmill belt to the center of mass and **v**^COM^ is the center of mass velocity.

Computed in the slow belt frame, the pre- and post-collision angular momenta are

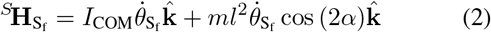

and

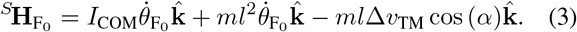

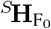 contains Δv_TM_ because the post-collision velocity includes the belt speed difference when expressed in the slow frame.

Setting Equations (2) and (3) equal and solving for the angular velocity post-collision yields:

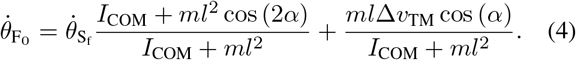

Note that during the fast belt collision, the belt speed difference increases the wheel’s angular velocity.

#### 2.3.3 Rotation on the Fast Belt

The fast belt rotation is similar to the slow belt rotation, and an energy balance in the fast belt reference frame can be used to compute the velocity at the end of the rotation given the velocity at the beginning. The angular velocity after the fast belt rotation 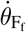, is

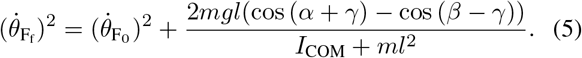

#### 2.3.4 Collision onto the Slow Belt

The slow belt collision is similar to the fast belt collision except that it is the velocity before the collision, rather than after, that must be transferred to the slow belt frame when computing the conservation of angular momentum:

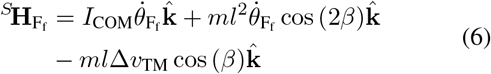

and

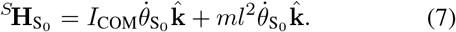

Combining Equations (6) and (7) yields the governing equation for the slow belt collision:

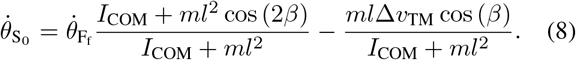

During the slow belt collision, the belt speed difference decreases, rather than increases, the wheel’s angular velocity.

To solve for the fixed point 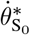 that enables steady walking, Equations (1), (4), (5), and (8) can be combined to create a single equation.

### 2.4 Governing Equations for Rotation

Whereas energy balance and conservation of momentum can be used to identify fixed points and fully characterize collisions, numerical integration is required to calculate how the split-belt wheel moves during the rotations. This allows calculation of the time spent in each rotation and of the forces acting on the wheel throughout the cycle.

For each rotation phase, we have:

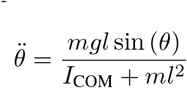

where *θ* is the angle from vertical 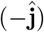 to the contacting spoke, positive in the 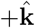 direction (Fig. 2). We obtain the state trajectory by numerical integration. We then calculate the horizontal and vertical accelerations and use them to calculate the ground reaction forces. The forces are used to calculate the work from the treadmill on the wheel and to exclude cases where the wheel loses contact with the ground.

### 2.5 Slow-Belt Walking Speed

Comparing the wheel’s walking speed across split-belt conditions requires a careful definition of “walking speed,” just as human split-belt walking speed is not as simple as averaging the two belt speeds. In this paper, “slow-belt walking speed” refers to the amount the wheel’s center of mass moves forward in the slow belt frame during one gait cycle divided by the total time for that cycle.

During the slow belt rotation, the forward movement of the center of mass is

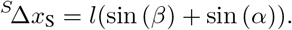

During the fast belt rotation, the contact point *P* moves backward relative to the slow belt, and the net forward movement is

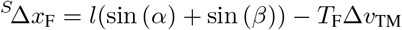

where *T_F_* is the duration of the fast belt rotation. The wheel’s slow-belt walking speed is therefore

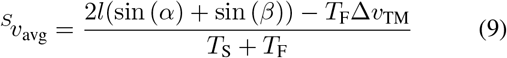

where *T*_S_ is the duration of the slow belt rotation.

The slow-belt walking speed can be transferred to the stationary lab frame by subtracting the slow belt velocity. The slow-belt walking speed can be thought of as the wheel’s forward velocity in the lab frame if the slow belt is stationary, or as the slow belt velocity necessary for the wheel to remain stationary in the lab frame while walking on the treadmill (the station-keeping speed).

### 2.6 Energy and Work during Rotations

We calculated velocities, energy, power, and work during the rotations to understand how energy is transferred between the treadmill and the split-belt rimless wheel. We performed all calculations in the slow belt reference frame.

#### 2.6.1 Rotation on the Slow Belt

During the slow belt rotation, the wheel’s kinetic energy is

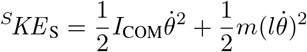

and the potential energy is

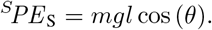

In the slow-belt frame, total energy remains constant and no work is done during the slow belt rotation because there is no relative movement of the contact point *P*.

#### 2.6.2 Rotation on the Fast Belt

Velocity, energy, power, and work in the slow belt frame during the fast belt rotation are affected by the belt speed difference Δν_TM_. During the fast belt rotation, the velocity of the center of mass expressed in the slow belt frame is

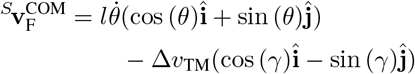

where *γ* is the treadmill incline and Δν_TM_ has been subtracted out to transfer the velocity into the slow belt frame. The wheel’s kinetic energy is then

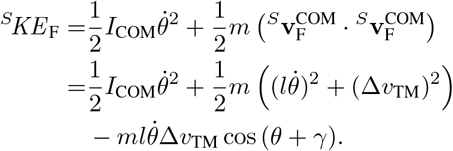

The wheel’s potential energy in the slow belt reference frame during the fast belt rotation is

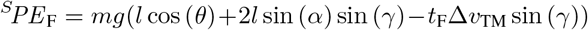

where *t_F_* is the time since the fast belt collision. The 2*l* sin (*α*) sin (*γ*) term accounts for the difference in height of the contact point *P* before and after the fast belt collision, and the –*t*_F_Δν_pM_ sin (*γ*) term accounts for the downward movement of the contact point during the fast belt rotation (Appendix C).

The treadmill can perform work on the wheel during the fast belt rotation because there is relative motion at the contact point *P* and a force **F** acting between the treadmill and the wheel. The velocity of P during the fast belt rotation, viewed from the slow frame, is

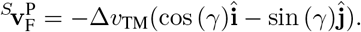

The work ^S^W_F_ of the treadmill on the wheel is

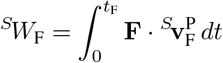

where *t*_F_ is the time since the fast belt collision. The force **F**, the ground reaction force acting on the wheel at the contact point, depends on the wheel’s angular position, velocity, and acceleration.

### 2.7 Energy and Work during Collisions

We used the pre- and post-collision velocities to compute the wheel’s kinetic energy before and after each collision, subtracting those to find the effect of the collision. Potential energy does not change in the collision because the center of mass does not move in the instant of the collision. We completed all calculations in the slow belt reference frame.

The energy change in the fast belt collision is the difference between the post-collision kinetic energy and the pre-collision kinetic energy:

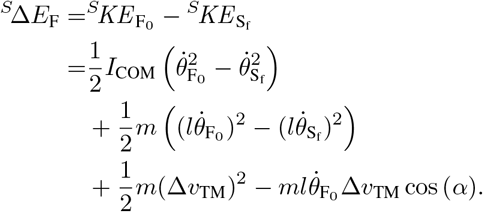

For the slow belt collision, we have:

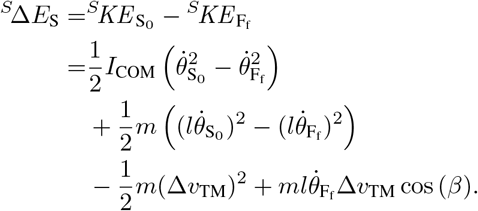

For a steady gait cycle on level ground, the total energy change across an entire gait cycle is zero, meaning

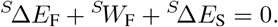

For a steady gait cycle uphill, the total energy change equals the change in the wheel’s overall potential energy from moving uphill (Appendix C).

### 2.8 Stability and Disturbance Rejection

We used two different methods to evaluate the split-belt rimless wheel’s stability and disturbance rejection. We used eigenvalue analysis to understand local stability in a narrow region around the limit cycle, and we used variable terrain to approximate real-world conditions and understand the wheel’s ability to reject repeated disturbances over many steps.

#### 2.8.1 Eigenvalue Stability Analysis

We characterized the wheel’s response to small angular velocity perturbations using a finite differencing approach to estimate the eigenvalue of our one-dimensional system as

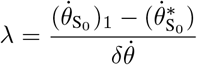

where 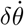 is the small offset in angular velocity added at the beginning of the slow belt rotation, 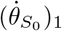 is the angular velocity after one complete gait cycle, and 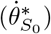 is the fixed point angular velocity for the beginning of the slow belt rotation. In a small region around the fixed point, λ is the same regardless of the size of 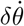 or where in the cycle 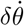 is introduced. Eigenvalues with magnitude less than one indicate asymptotic stability, and positive eigenvalues less than one indicate that the wheel gradually returns to the fixed point without overshoot.

#### 2.8.2 Variable Terrain Disturbance Rejection

Eigenvalue analysis has limited relevance in the presence of large disturbances. To capture the cumulative effects of repeated disturbances and move outside the small region where linear analysis is valid, we simulated the wheel walking over uneven terrain (Fig. 3) where the ground height changed before every collision between a spoke and the ground (Kim and Collins 2017). Before each variable terrain simulation, we generated 1000 ground height values, evenly and randomly distributed between two limits (±GH). We quantified stability as the maximum value of GH for which the split-belt rimless wheel could walk 500 cycles (1000 steps) without failure.

**Figure 3.**
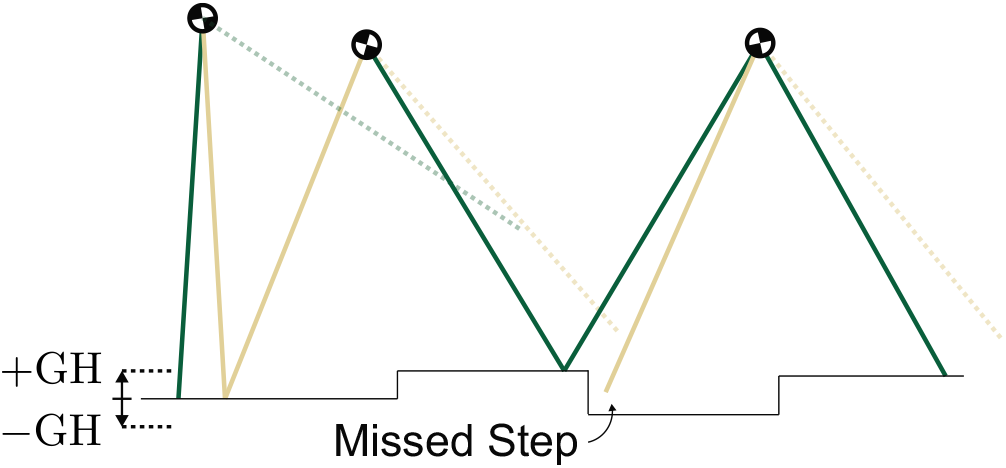
A sample variable terrain with ground heights distributed between +GH and −GH. The wheel’s three failure modes were missing a step, failing to reach the top during a rotation, and entering a flight phase.

Failure could occur in one of three ways. First, the wheel could slow down and fall backward, failing to reach vertical after a step up. Second, the wheel could contact the same belt twice in a row, missing a step onto the other belt. Finally, the wheel could lose contact with the treadmill altogether, rotating so quickly as to enter a flight phase.

### 2.9 Nondimensionalization

The input parameters and outcome measures of the model can be nondimensionalized with respect to the wheel’s leg length *l*, mass *m*, and gravitational constant *g*. When calculating the moment of inertia *I*_COM_, we assume that the radius of gyration *r*_gyr_ scales with *l*. Speed terms, such as belt speed difference and slow-belt walking speed, are nondimensionalized through division by 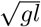. The ground height disturbance limits ±GH are nondimensionalized through division by *l*. Mass is eliminated by dividing the energy by *m*.

## 3 Simulation Results and Discussion

In this section, we use four model variations to discuss the impact of changing the wheel’s collision angles and moment of inertia, as well as the belt speed difference and treadmill incline.

### 3.1 Mass and Leg Length Do Not Affect Nondimensionalized Outcomes

When results are nondimensionalized, the wheel’s mass and leg length have no effect on outcomes. Mass simplifies out of all equations that govern the wheel’s motion, and system energy scales linearly with mass. Similarly, leg length does not alter nondimensionalized speed, stability, or energy. To facilitate quick comparisons between split-belt rimless wheel walking and human split-belt treadmill walking, we present the simulation results for a leg length of one meter rather than presenting them in dimensionless form.

### 3.2 Model Variation 1: The Simplest Split-Belt Rimless Wheel

#### 3.2.1 Assumptions and Description of Motion

We began with the simplest possible version of the split-belt rimless wheel. We set the mass moment of inertia and the treadmill incline angle to zero. By setting the angular offset between the two individual rimless wheels to be infinitesimally small, we achieved a slow belt collision angle β of approximately zero.

These parameter choices simplify the dynamics of the split-belt rimless wheel (Fig. 4A). Because *β* is infinitesimally close to zero, the wheel begins each slow belt rotation with the spoke vertical. As it rotates down through the angle *α*, the center of mass’s vertical motion is exclusively downward. In the fast belt rotation, as the wheel rotates up through *α*, the center of mass’s vertical motion is exclusively upward. Because *I*_COM_ is zero, the ground reaction forces and collision impulses must always act along the contacting leg; if not, the resulting angular acceleration would be infinitely large. In the slow belt collision, no energy is lost because the infinitesimally small collision angle means the center of mass velocity does not need to be redirected.

**Figure 4.**
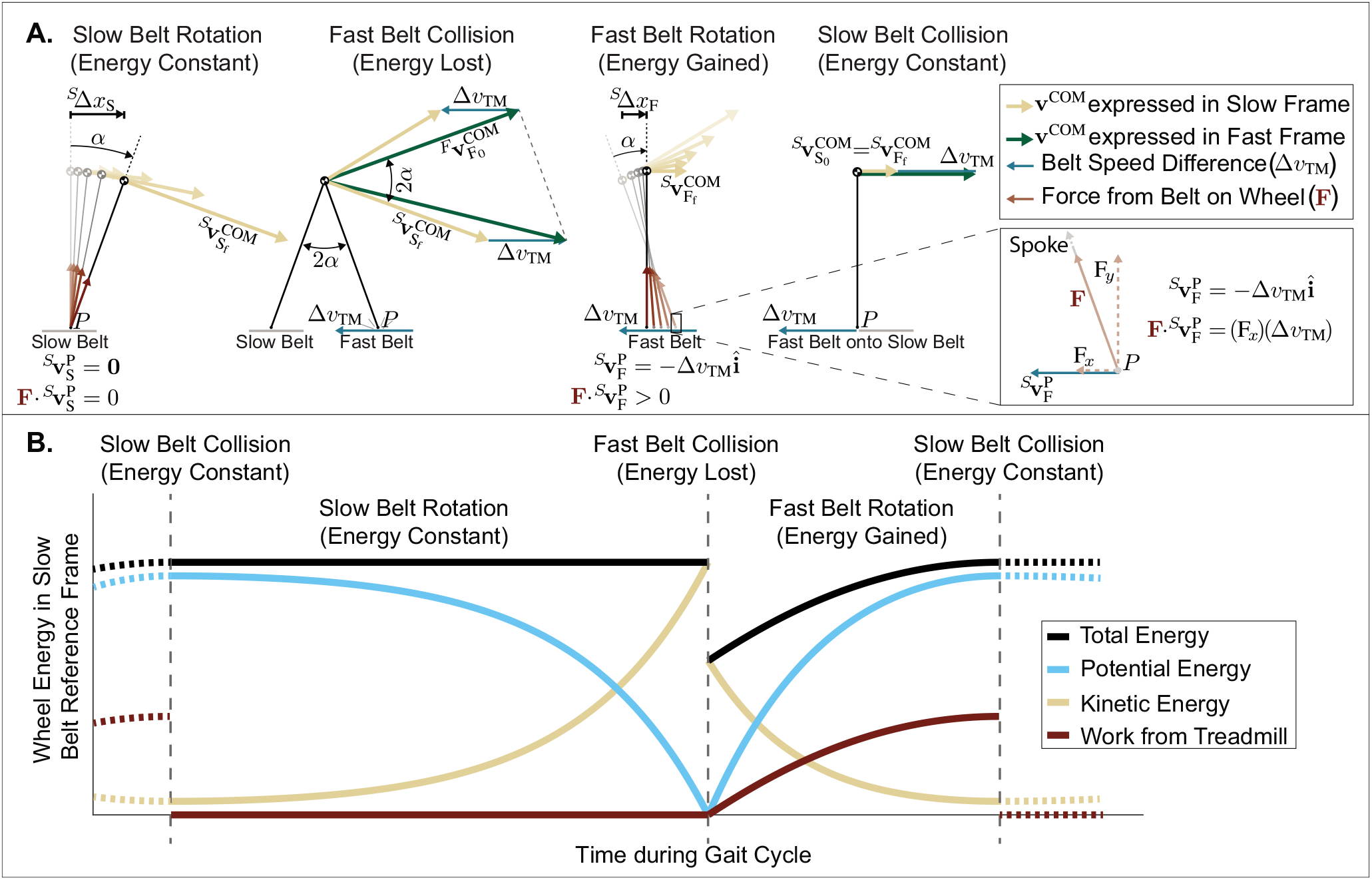
**A**. Hodographs for a sample steady gait cycle of the simplest split-belt rimless wheel with slow belt collision angle of zero. Hodographs are drawn in the slow belt reference frame, so the contact point *P* is stationary during the slow belt rotation. The center of mass velocity **v**^COM^ is transferred between the slow and fast belt frames with the addition of Δ*ν*_TM_. To visualize the fast belt collision, the pre-collision velocity is transferred to the fast frame since the collision occurs on the fast belt, but then post-collision velocity is transferred back to enable an energy comparison in the slow belt frame. **B**. Plot tracking the split-belt rimless wheel’s energy in the slow belt reference frame through a steady cycle. During the slow belt collision and the slow belt rotation, the wheel’s total energy remains constant. The wheel loses energy in the fast belt collision but recovers that energy in the fast belt rotation such that total energy at the end of the cycle equals the total energy at the beginning of the cycle.

#### 3.2.2 Passive Steady Walking

When walking steadily, the wheel starts and ends each gait cycle with the same energy (computed in the slow frame in Fig. 4B). The wheel begins the cycle with a slow belt rotation during which the contact point remains stationary and the treadmill does no work as the wheel evenly exchanges potential and kinetic energy. During the fast belt collision, the wheel loses energy as the center of mass velocity is redirected. The wheel recovers that energy during the fast belt rotation because the treadmill does net positive work on the wheel. Finally, the energy remains constant during the vertical slow belt collision because the center of mass velocity requires no redirection.

The split-belt treadmill injects energy during the fast belt rotation because the contact point P moves backwards while there is a backwards component to the ground reaction force **F** (Fig. 4A Inset). This alignment between **F** and the contact point velocity 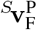 means the power 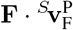 from the treadmill on the wheel is positive throughout the entire fast belt rotation, making the net work positive. The ground reaction force has a backward horizontal component because *P* is in front of the center of mass for the entire rotation, so the force which points along the leg from *P* toward the center of mass has a backwards component.

Another way to understand the energy gain during the fast belt rotation is that the treadmill helps lift the center of mass. If the treadmill was not moving, the raising of the center of mass during the rotation would be accomplished solely through an exchange of potential and kinetic energy. When the fast treadmill is moving, the belt pulls the bottom end of the spoke backward, which forces the spoke underneath the wheel center and helps lift up the center of mass. The treadmill does some of the work to lift the center of mass and increase the wheel’s potential energy, allowing the wheel to keep more of its kinetic energy instead of having to trade it for potential energy. By the end of the fast belt rotation, because of the help of the treadmill, the wheel has more total energy than when it started the fast belt rotation.

The net positive work during the fast belt rotation is possible because of two important asymmetries. The belt speed difference gives the contact point *P* a non-zero velocity in the slow belt reference frame, enabling non-zero power during the fast belt rotation. Without an asymmetry in belt speeds, no work can be done because the contact point would always be stationary in both belt reference frames. The asymmetry in collision angles, and the resulting longer step onto the fast belt and infinitesimally small step onto the slow belt, mean the wheel only rotates up during the fast belt rotation, instead of evenly rotating both up and down as it would if the angles were equal. This keeps the power positive by allowing the contact point *P* to remain in front of the center of mass during the entire fast belt rotation, meaning the horizontal component of force stays aligned with the contact point velocity.

When all else is equal, the wheel extracts more energy from the treadmill when it spends longer in the fast belt rotation. A slower rotation means having similar power applied for a longer period of time. The force is primarily angle-dependent, but the contact point displacement increases with the duration of fast-belt stance, increasing the total product of force and displacement. Rotating more slowly also means the wheel enters the fast belt collision more slowly and loses less energy in the collision. The rotation duration is inversely proportional to the angular velocity because the spokes are a fixed distance apart, so a longer rotation duration requires lower angular velocity.

#### 3.2.3 Feasibility Region

There exists a range of belt speed difference Δ*ν*_TM_ and fast belt collision angle 2*α* combinations for which the simplest split-belt rimless wheel can walk steadily on level ground, instead of slowing to a stop like a regular rimless wheel (Fig. 5). The feasible region is bounded at the y-axis because as *α* approaches zero, the wheel becomes a regular rimmed wheel that can roll steadily on level ground for any initial conditions. Large values of *α* are infeasible because the wheel loses more energy in the fast belt collision than it can recover during the fast belt rotation. Large values of Δ*ν*_TM_ are infeasible because, in order to balance the collision loss and rotation gain, the wheel must rotate so quickly that it loses contact with the treadmill.

**Figure 5.**
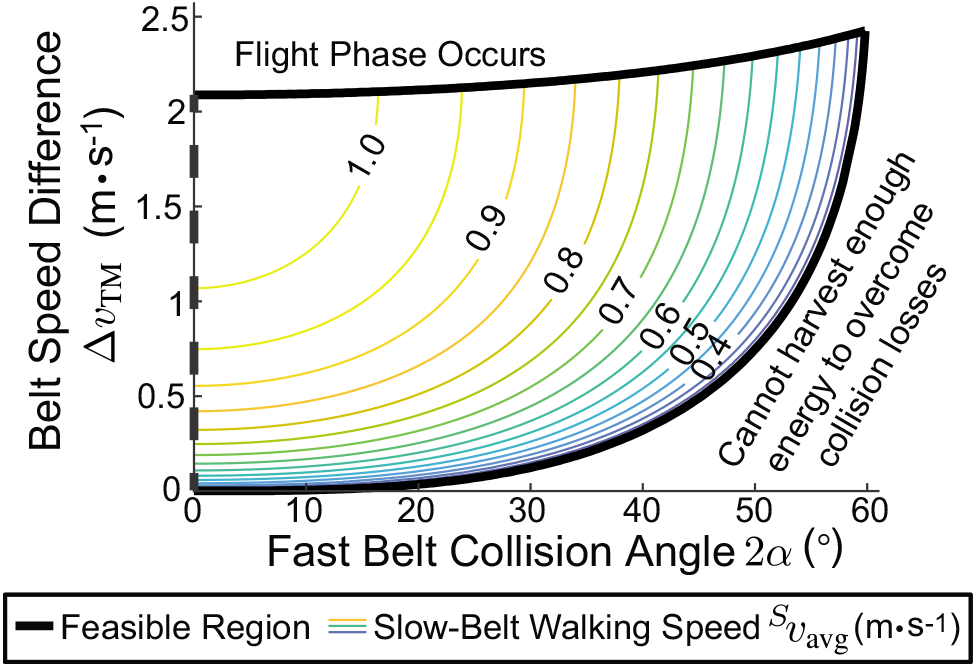
The simplest split-belt rimless wheel, with a slow belt collision angle of zero, can walk steadily for a range of fast belt collision angles and treadmill belt speed differences. For all combinations of 2*α* and Δ*ν*_TM_ within the feasible region, the wheel has a positive slow-belt walking speed 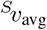. Increasing Δ*ν*_TM_ increases 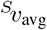, while increasing *α* decreases 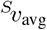.

#### 3.2.4 Effects of Belt Speeds and Collision Angles

The simplest split-belt rimless wheel’s slow-belt walking speed increases both when the belt speed difference increases and when the fast belt collision angle decreases (Fig. 5). Increasing Δ*ν*_TM_ decreases energy lost in the fast belt collision and increases energy gained in the fast belt rotation (Appendix D). When *α* is held constant but Δ*ν*_TM_ increases, the angular velocity must increase or the rotation gain would be larger than the collision loss. The increase in angular velocity results in an increased slow-belt forward walking speed because, although the fast treadmill moves the wheel farther back during the fast belt rotation, the rotation durations in Equation (9) decrease enough that the overall walking speed increases (Appendix E). Decreasing *α* decreases both the energy lost in collision and the energy gained in rotation, but the collision loss decreases more than the rotation gain (Appendix D). When *α* decreases, the angular velocity must increase to balance the energy, resulting in an increased overall slow-belt walking speed (Appendix E). If the slow belt is stationary, then the increased slow-belt walking speed corresponds to the wheel moving forward faster in the lab frame. If the wheel is to maintain constant average position in the lab frame, then the increased walking speed means the slow treadmill belt must be set to a higher speed.

#### 3.2.5 Stability

The same dynamics that enable the simplest split-belt rimless wheel to walk steadily make it asymptotically stable to speed perturbations. The eigenvalue for the discrete, linearized stride-to-stride state transition matrix is always positive and less than or equal to one, meaning the wheel behaves like an overdamped system as it returns to its steady speed following a perturbation (Appendix F). If the wheel is perturbed to rotate more slowly than its steady speed, it loses less energy in the fast belt collision and spends more time on the fast treadmill belt, gaining more energy in the rotation than it had lost in the collision. The wheel ends the cycle with higher energy and continues speeding up with each cycle until returning to its steady rolling speed. The opposite effect occurs if it is perturbed to roll too quickly.

Although this simple version of the split-belt rimless wheel with an infinitesimally small slow belt collision angle *β* tolerates perturbations in angular velocity, it has no capacity to reject ground height disturbances. An infinitesimal difference between the heights of the slow and fast belts results in inappropriate foot sequencing with the wheel contacting the same belt twice in a row. A more complex wheel with a non-zero slow belt collision angle is required to understand the wheel’s ground height disturbance tolerance.

### 3.3 Model Variation 2: The Angular Offset Split-Belt Rimless Wheel

#### 3.3.1 Assumptions and Description of Motion

We generalized the simplest rimless wheel by removing the assumption of infinitesimally small angular offset between the two individual rimless wheels, thereby allowing the slow belt collision angle *β* to be non-zero. We kept the mass moment of inertia and the treadmill incline angle at zero.

Because of these assumptions, the ground reaction forces and impulses still point along the contacting leg throughout the cycle, but the center of mass’s vertical motion within each rotation is no longer monotonic (Fig. 6A). The center of mass rotates up through *β* and down through *α* in the slow belt rotation, and up through *α* and down through *β* in the fast belt rotation. Assuming *α* > *β*, the center of mass height decreases overall in the slow belt rotation and increases in the fast belt rotation. Additionally, the slow belt collision no longer occurs at vertical, and the center of mass velocity must be redirected before the wheel can begin its next slow belt rotation.

**Figure 6.**
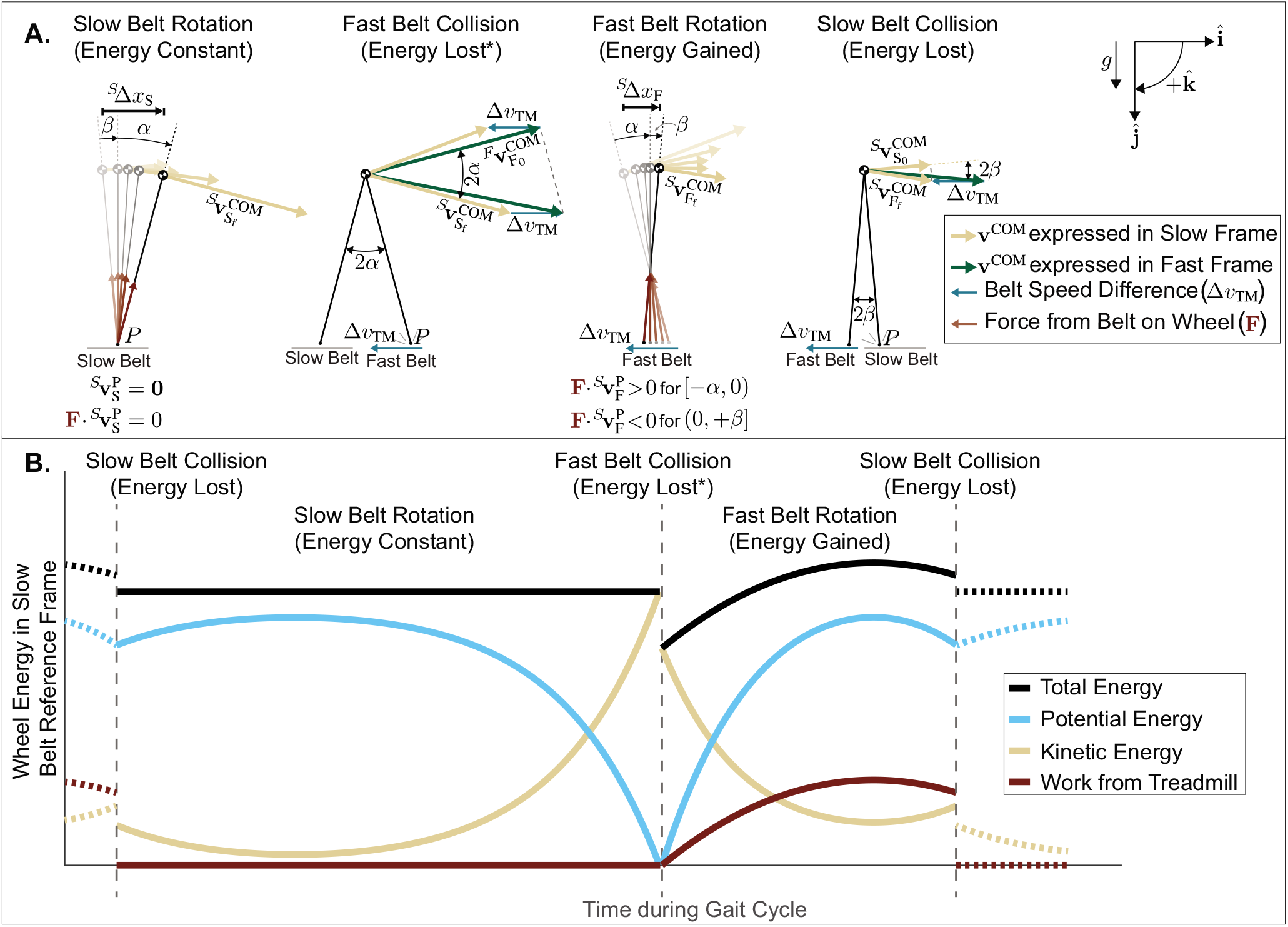
**A**. Hodographs of a sample steady gait cycle for a rimless wheel with non-zero angular offset (2*β* > 0). The pre-collision velocities must be transferred from the trailing leg’s reference frame to the leading leg’s reference frame to enable visualization of the collision impulse. During the rotations, the center of mass moves both up and down, and the work from the treadmill during the fast belt rotation is positive as the wheel rises up through *α* and negative as it falls down through *β*. **B**. Plot tracking the wheel’s energy in the slow belt reference frame through the cycle. The wheel typically loses energy in the fast belt collision*, but the treadmill does net positive work on the wheel during the fast belt rotation to recover the lost energy. The energy lost in the slow belt collision brings the wheel’s total energy back to its value from the beginning of the cycle. The magnitude of the slow belt collision loss has been exaggerated for clarity. *Cases exist for high Δ*ν*_TM_ where energy is gained, not lost, in the fast belt collision (Appendix D).

#### 3.3.2 Passive Steady Walking with Non-zero β

The splitbelt rimless wheel with a non-zero angular offset is able to walk steadily through the same mechanism as in the simplest case. Viewed in the slow belt reference frame, the wheel gains enough energy in the fast belt rotation to overcome the energy lost in collisions, but the non-zero *β* does cause a few differences (Fig. 6B). The slow belt collision results in energy loss because of the velocity redirection of the center of mass. The fast belt collision sometimes results in energy gain instead of energy loss when Δ*ν*_TM_ is high (Appendix D). Finally, the fast belt rotation always results in energy gain, but the amount of energy gained decreases as *β* increases.

Increasing *β* decreases the amount of net positive work done on the wheel during the fast belt rotation because the power is negative once the wheel passes through vertical. As the wheel rotates down through *β*, the horizontal component of the ground reaction force is opposite the direction of the contact point velocity. Another way to understand the adverse impact of *β* is that once the wheel passes vertical, the treadmill begins pulling the center of mass down, rather than helping lift it up. During the descent, the treadmill pulls the spoke out from under the wheel, stealing the wheel’s potential energy without allowing the wheel to evenly trade that potential energy for kinetic energy. For the net work to be positive during the fast belt rotation, step length asymmetry is required: *α* must be larger than *β* so that the wheel spends more time rising, with the treadmill doing positive work, than falling, with the treadmill doing negative work.

#### 3.3.3 Effect of β on Walking Speed and Feasible Region

Increasing the slow belt collision angle requires the wheel to walk more slowly (Fig. 7). The additional collision loss from the slow belt collision coupled with the decrease in net positive work during the fast belt rotation means the wheel must roll more slowly to decrease its collision costs and increase the energy absorbed during the fast belt rotation.

**Figure 7.**
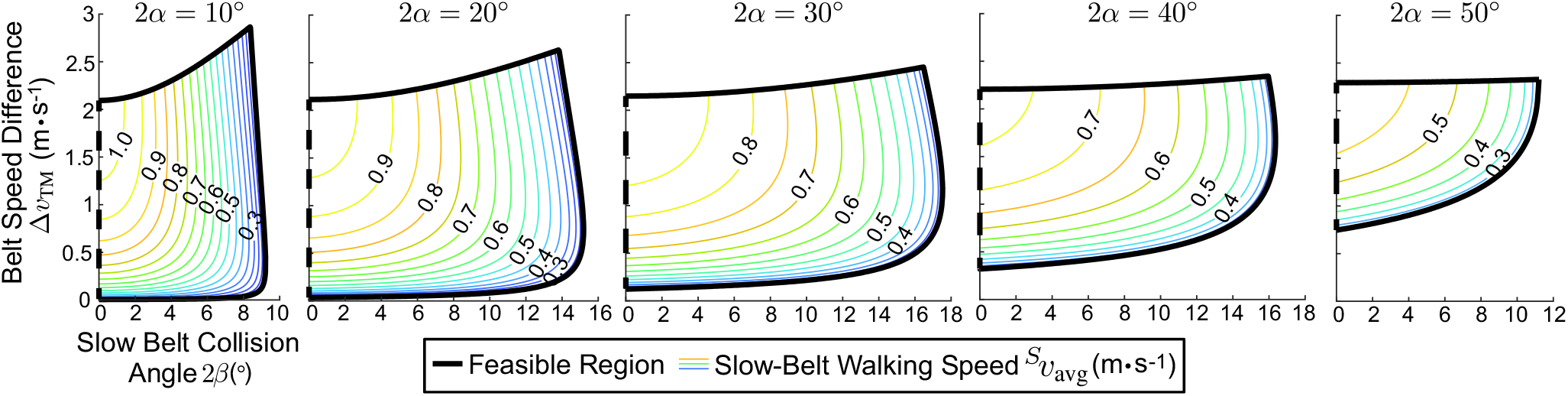
The split-belt rimless wheel can walk steadily for a range of belt speed differences and fast and slow collision angles. The largest range of slow belt collision angles is achieved when the fast belt collision angle 2*α* is intermediate. Increasing *β* decreases the wheel’s slow-belt walking speed 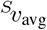 while increasing Δ*ν*_TM_ tends to increase 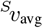.

The maximum feasible slow belt collision angle occurs with an intermediate fast belt collision angle (Fig. 7). If *α* is small, then the margin of (*α* − *β*), the limiting factor on how much work can be gained during the fast belt rotation, is small and quickly disappears as *β* increases. At the other extreme, when *α* is large, the wheel is barely able to overcome the fast belt collision loss even when *β* is zero and there is no energy lost in the slow belt collision.

#### 3.3.4 Effect of Belt Speed for Non-zero β

Increasing the belt speed difference tends to increase the wheel’s slow-belt walking speed even when *β* is non-zero (Figure 7). Despite the fact that increasing Δ*ν*_TM_ increases the slow belt collision loss (Appendix D), the benefits of higher Δ*ν*_TM_ outweigh the detriment. The fast belt collision loss decreases by more than the slow belt collision loss increases, and the higher Δ*ν*_TM_ increases the energy gained during the fast belt rotation. Together, those changes lead to faster steady-state angular velocities, which tend to result in higher steady walking speeds (Appendix E).

#### 3.3.5 Stability and Disturbance Rejection

The split-belt rimless wheel with non-zero *β*, like the simplest splitbelt rimless wheel, is asymptotically stable in response to small angular velocity perturbations. The eigenvalue for the discrete, linearized, stride-to-stride state transition matrix remains between zero and one for the feasible region of collision angles and belt speed differences.

The split-belt rimless wheel with non-zero *β* is also able to reject disturbances in ground height and walk over variable terrain. To understand the impact of each parameter, we chose a set of parameters well within the feasible region (2*α* = 30°, 2*β* = 8°, and Δ*ν*_TM_ = 1.0 m · s^−1^) and varied each parameter independently while leaving the other two parameters fixed (Fig. 8).

**Figure 8.**
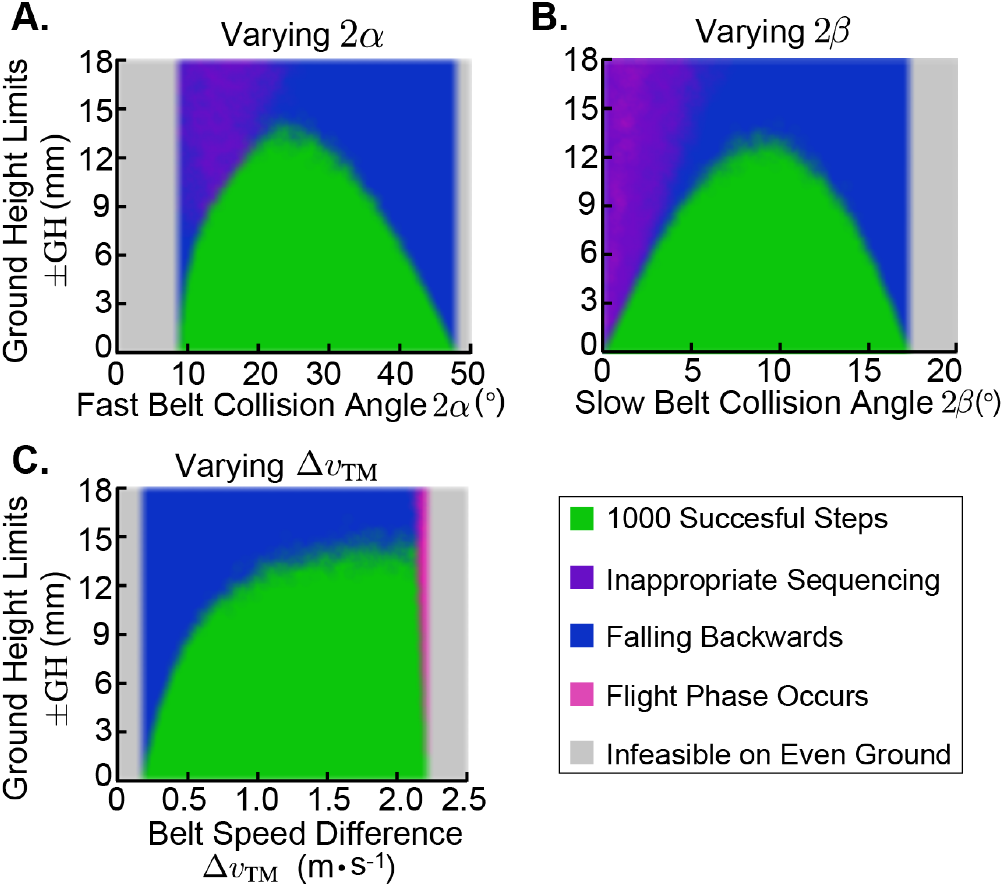
We varied the fast belt collision angle (**A**), the slow belt collision angle (**B**), and the belt speed difference (**C**) while holding the other parameters constant at 2*α* = 30°, 2*β* = 8°, and Δ*ν*_TM_ = 1.0 m · s^−1^. Each colored mark indicates the outcome of a variable terrain simulation for a given set of ground height limits ±GH. The split-belt rimless wheel with intermediate angles walking on a treadmill with high belt speed difference had the highest disturbance tolerance.

The wheel has maximal ground height disturbance tolerance when its collision angles are intermediate. With small collision angles, the wheel primarily fails due to inappropriate foot sequencing (purple region) or an inability to reach the top of a rotation following a step up (blue region). A disturbance seems larger relative to the step length when the collision angles are small (Appendix G), so a given disturbance size more significantly affects a wheel with small angles. A step up onto the slow belt can mean an early collision with the slow belt that drastically decreases the amount of time spent on the fast belt and the energy gained from the fast belt rotation. A wheel with large collision angles tends to fail by falling backward following a step up. On a uniform surface, a wheel with large collision angles barely completes its rotations and has very little margin to overcome disturbances that slow it down.

High belt speed difference enables the wheel to better tolerate ground height disturbances, as long as the wheel does not enter a flight phase (pink region). The wheel is able to sustain higher walking speeds with higher Δ*ν*_TM_, enabling a larger margin between the wheel’s average angular velocity and the minimum angular velocity needed to complete a given rotation. When Δ*v*_TM_ is high, the speed reduction accompanying a step up is less likely to prevent the wheel from completing its subsequent rotation. Belt speed difference appears beneficial both for ground height disturbance tolerance and for steady walking speed; holding the slow belt speed constant and making the fast belt move backward faster enables the wheel to walk forward more quickly and robustly.

### 3.4 Model Variation 3: Uphill Walking with the Simplest Split-Belt Rimless Wheel

#### 3.4.1 Assumptions and Description of Motion

While walking uphill, the split-belt rimless wheel can gain net positive energy by maintaining constant average kinetic energy and increasing gravitational potential energy with each cycle. For uphill walking of the simplest split-belt rimless wheel (*β* = *I*_COM_ = 0), the wheel’s motion and energy through a cycle are very similar to the previous cases apart from a few key exceptions. First, in the slow belt frame, the center of mass ends the cycle higher than its starting position, meaning the energy gained during the fast belt rotation must compensate for both the energy lost in collision and the overall increase in wheel height. Second, during the slow belt rotation, the contacting spoke begins normal to the incline instead of vertical, so the slow belt rotation includes a few degrees of upward rotation. Finally, during the fast belt rotation, the entire wheel moves backward and down relative to the slow belt, so the change in the wheel’s potential energy is dependent on the duration *T_F_* of the fast belt rotation. A slower rotation results in a lower overall potential energy increase for the cycle because the wheel spends more time moving down.

#### 3.4.2 Incline Effect on Feasibility and Walking Speed

We found that the simplest split-belt rimless wheel could successfully harness enough energy from the treadmill to walk steadily up surfaces with a small incline angle, *γ*. As *γ* increased, the wheel walked more slowly for a given set of collision angles and belt speed difference, and the size of the feasible region decreased (Fig. 9). Although not shown, feasible solutions existed for small values of *γ* even when the slow belt collision angle was non-zero.

**Figure 9.**
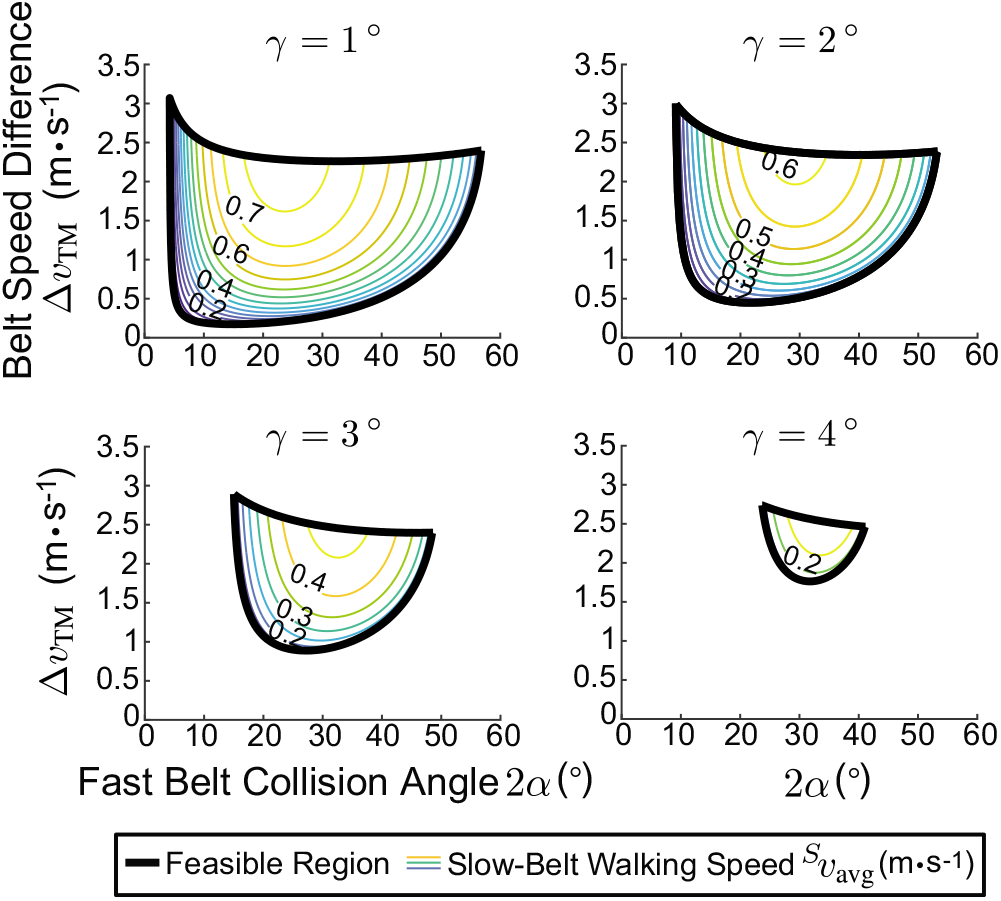
The split-belt rimless wheel can walk steadily uphill, but the feasible region decreases as the treadmill incline *γ* increases. Shown here are solution spaces for an infinitesimally small slow belt collision angle (*β* = 0), but solutions do exist when *β* is non-zero. The wheel walks fastest and can walk up the highest inclines when the fast belt collision angle is intermediate. Increasing the belt speed difference also increases the wheel’s slow-belt walking speed.

By walking more slowly for larger inclines, the wheel could extract enough energy during the fast belt rotation to achieve the overall increase in potential energy at the end of the cycle associated with moving uphill. Slower speed meant more time in the fast belt rotation extracting work from the treadmill and a lower total potential energy gain required per cycle. The need for a slower speed for the same fast belt collision angle, however, meant that larger values of *α* became infeasible at higher inclines. A wheel with large *α* cannot slow down enough to adequately increase its energy capture without falling backward during rotations.

There is also a minimum *α* for uphill walking that increases with *γ*, a contrast to level ground walking where *α* can approach zero. As the incline increases, driving up the energy that must be gained per cycle, the minimum *α* increases because wheels with small angles cannot gain enough energy from the treadmill during the fast belt rotation to achieve the necessary net energy increase. An ideal rimmed wheel can roll steadily on level ground, losing no energy and gaining no energy, but it cannot increase its energy over time by rolling steadily uphill.

The impact of increasing the belt speed difference Δ*ν*_TM_ is similar for uphill walking and level ground walking. A larger Δ*ν*_TM_ allows the wheel to walk with higher speeds and up larger inclines because the belt speed difference allows for more energy capture within a cycle.

### 3.5 Model Variation 4: The Split-Belt Rimless Wheel with Non-Zero Rotational Inertia

#### 3.5.1 Assumptions and Description of Motion

To verify that these behaviors can be achieved with physically realistic mass distributions, we conducted a final set of simulations while varying the radius of gyration *r*_gyr_ between zero and the spoke length *l*. We kept the slow belt collision angle infinitesimally small and set the treadmill incline at zero. Non-zero *I*_COM_ means the ground reaction forces and collision impulses are no longer guaranteed to be directed along the contacting leg, and the vertical slow belt collision is no longer lossless. Because the wheel has both linear and rotational kinetic energy, angular velocity changes less during a rotation and the wheel can begin a rotation more slowly but still reach the top. The fast belt collision is less costly with rotational inertia, but less energy is gained in a fast belt rotation that begins with identical angular velocity.

#### 3.5.2 Inertia Effect on Feasibility, Walking Speed, and Disturbance Rejection

The model generally behaves similarly to the no-inertia case, but we found that as the rotational inertia increased, the wheel walked more slowly for a given set of collision angles and belt speed difference, and the feasible region increased (Fig. 10A, B). Lower fast belt rotation gains and the cost of the slow belt collision dominated the lower fast belt collision costs, leading to a slower steady walking speed. Higher collision angles became feasible because the wheel could rotate more slowly while still completing rotations, and higher belt speed differences were feasible because the wheel could balance its cycle energy without increasing its angular velocity so much as to enter a flight phase.

**Figure 10.**
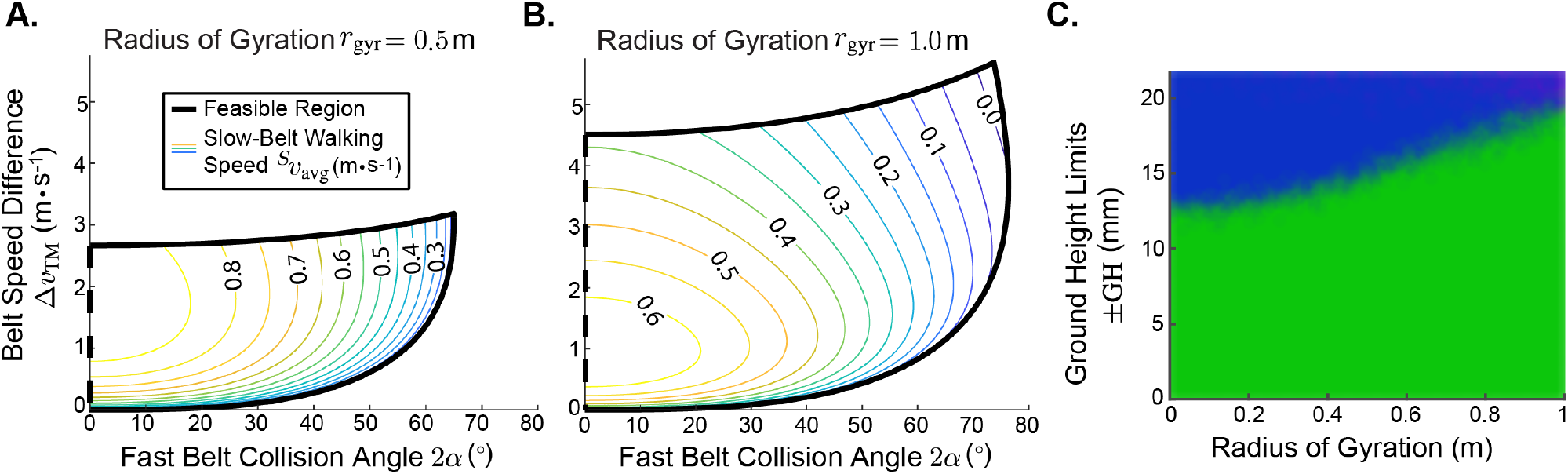
When the slow belt collision angle is infinitesimally small, as the inertia increases from *r*_gyr_ = 0.5 m (**A**) to *r*_gyr_ = 1.0 m (**B**), the split-belt rimless wheel walks more slowly but the feasible solution region increases. We varied the radius of gyration *r*_gyr_ while holding the fast belt collision angle 2*α* at 30°, the slow belt collision angle 2*β* at 8°, and the belt speed difference Δ*ν*_TM_ at 1.0 m · s^−1^(**C**). Each colored mark indicates the outcome of a variable terrain simulation for a given set of ground height limits ±GH. Green indicates successful completion of 1000 steps, blue indicates a failure from falling backward, and purple indicates a failure from inappropriate step sequencing. Increasing the rotational inertia increased the wheel’s disturbance rejection.

We also found that increasing the rotational inertia increases the wheel’s tolerance to ground height disturbance for a representative fast belt collision angle 2*α* = 30°, slow belt collision angle 2*β* = 8°, and belt speed difference Δ*ν*_TM_ = 1.0 m · s^−1^ (Fig. 10C). Having rotational kinetic energy means the relative energy loss associated with a step up of specific height is smaller relative to the wheel’s total energy.

### 3.6 Implications for Human Split-Belt Walking

Insights from the split-belt rimless wheel can help us understand human split-belt treadmill adaptation. During tied-belt walking, the treadmill cannot contribute any net work (Sánchez et al. 2019), but the treadmill can perform work during split-belt walking that allows the rimless wheel to walk steadily. For example, the wheel can station-keep on a treadmill with belt speeds of 0.5 and 1.5 m · s^−1^ without requiring any energy, but it could not walk steadily with both belts at 0.5 m · s^−1^. If human mechanics mirror wheel mechanics, this suggests that people might require less energy to walk on belts split at 0.5 and 1.5 than on belts tied at 0.5. There is a whole family of wheel solutions enabling energy-free station-keeping with these belt speeds (Fig. 11), indicating that many possible gaits exist for humans to take advantage of the treadmill.

**Figure 11.**
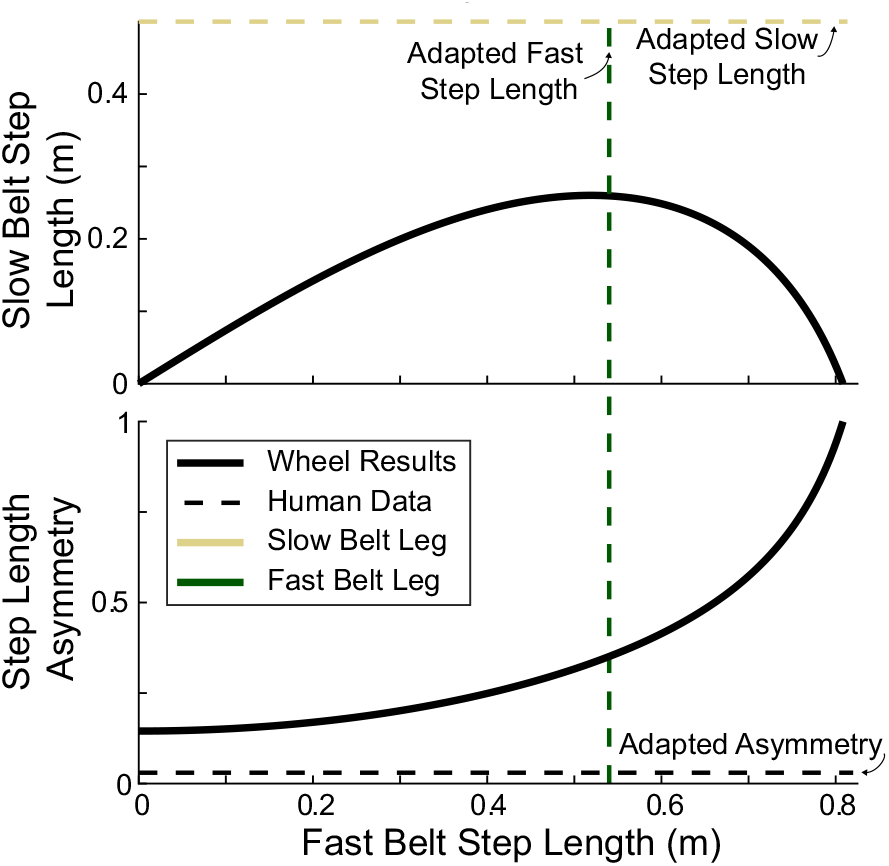
With treadmill speeds of 0.5 and 1.5 m · s^−1^, the split-belt rimless wheel can walk steadily while maintaining constant average position in the lab frame (see Extension 1). There is a range of step length combinations with associated step length asymmetry which all allow the wheel to station-keep. Human data for final adapted step lengths and step length asymmetry for this belt speed combination (Sánchez et al. 2020) are included for comparison between human and rimless wheel walking.

There is evidence that people adapt to the split-belt treadmill in a method consistent with taking advantage of treadmill work, but people do not adapt to the full extent possible. All configurations which allow the wheel to station-keep require positive step length asymmetry (Fig. 11), and previous work has shown that people gradually transition from negative to positive step length asymmetry when adapting to the split-belt treadmill (Sánchez et al. 2020). This evolution of step length asymmetry is associated with a decrease in energy cost between early and late adaptation, consistent with the theory that people are adapting to take advantage of positive work performed by the treadmill. People do not, however, adopt an asymmetry as extreme as that predicted by the model for their chosen fast belt step length, plateauing at just 0.03 after 45 minutes (Sánchez et al. 2020) rather than continuing to the model asymmetry of 0.35. The energy savings associated with human split-belt walking also fall well below the model prediction. We have not previously observed people achieving lower energy costs for walking on split belts than for walking on tied belts at the slow belt speed, despite the model’s indication that it might be possible. Instead, energy costs during split-belt walking are either even with or slightly below the cost of walking on tied belts at the average speed (Sánchez et al. 2019, 2020; Stenum and Choi 2020; Finley et al. 2013).

Humans adapt to a fast step length well within the family of model solutions, but their slow step length remains far longer than the model predicts would be optimal. To increase their step length asymmetry and better take advantage of the treadmill, people would need to decrease their slow belt step length. By comparing the gait adopted by humans to the model’s family of energy-free solutions, we can gain insight into which other aspects of split-belt walking could be interfering with people’s ability to take advantage of a belt speed difference to decrease their energy costs.

A high cost of swinging the legs could negate the benefits of certain gaits, but when we approximate swing costs as if the rimless wheel model had to swing its legs, the model gait does not encode significantly higher swing costs than the gait to which humans adapt. Previous work has indicated that swinging the legs at higher than their natural frequency can significantly increase the metabolic cost of walking (Doke et al. 2005), and swing costs trade off with push-off costs to control the optimal speed-step length relationship (Kuo 2001). Some of the model gaits with short steps onto both the fast and slow belts (left side of Fig. 12A) are unrealistic due to swing times that are shorter than observed even when people are instructed to walk with the highest step frequency possible (Nilsson and Thorstensson 1987). The other gaits, however, including the wheel solution at humans’ adapted fast belt step length, have swing times nearly equal to or longer than humans’ adapted step times, suggesting that those gaits are feasible. The average swing leg velocity (Fig. 12B), estimated as the leg’s total swing angle divided by the swing time, (*α* +*β*)/*T*, is lower for all the model gaits than for humans’ adapted gait, indicating that swing leg velocity is probably not preventing people from shortening their slow belt step length to adopt the model solution.

**Figure 12.**
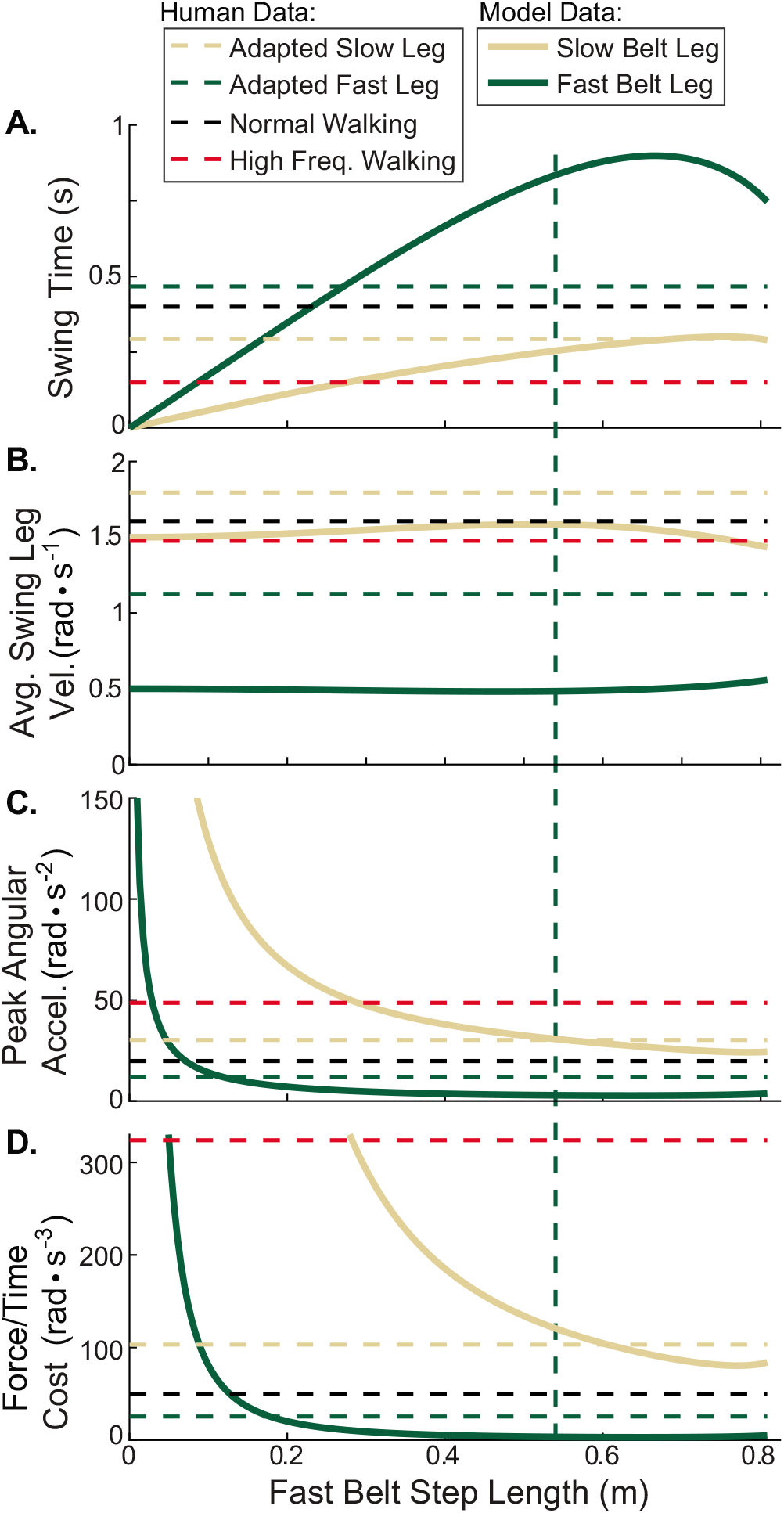
For the family of solutions in Fig. 11 enabling station-keeping on belts at 0.5 and 1.5 m · s^−1^, we found the swing time, average swing leg angular velocity, peak swing leg angular acceleration, and swing cost proportional to peak force divided by swing time. Using published human step length and swing time data for various types of walking, we consider whether the model gaits are feasible and whether they encode high swing costs. Comparison data include adapted split-belt walking at 0.5 and 1.5 m · s^−1^ (Sánchez et al. 2020), normal walking at 1.0 m · s^−1^, and walking with maximal step frequency at 1.0 m · s^−1^(Nilsson and Thorstensson 1987).

Previous work by Kuo (2001) and Doke et al. (2005) suggests that a more accurate metric for swing cost is peak force divided by the duration of swing; this metric does not, however, indicate a high swing penalty for all of the model’s gaits. Modelling the swing leg with pendular dynamics and a simplified trajectory *θ* = *A* cos (*ωt*), we can represent the peak force as proportional to the peak angular acceleration, *Aω*^2^ (Fig. 12C). We define the amplitude 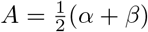 and the frequency 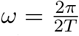 where *α* and *β* are the collision angles and *T* is the duration of swing. The force/time metric accounts for muscle inefficiencies when producing force in short bursts and sets swing cost proportional to the peak acceleration divided by the swing time (Fig. 12D). Although the peak acceleration and force/time cost are high for gaits with short steps onto both belts, at the fast belt step length adopted by humans the swing cost for the model’s slow belt step length is similar to the cost for human’s adapted slow belt step length. The costs associated with swinging the leg slower than its natural frequency are less well understood, and the model’s gait does have a long swing time for the fast belt leg. Overall, however, it seems probable that people could adopt the model gait without incurring significantly higher swing costs, suggesting that swing costs are not the primary factor limiting people’s ability to take advantage of the split-belt treadmill.

As a brief side note, it is important to consider that short steps do not always require stepping with high frequency. If, as in the left side of Fig. 11, both steps are short, then step frequency must be high to achieve moderate walking speeds. When asymmetric steps are allowed, however, a gait may contain a long step onto one belt and a short step onto the other while keeping the duration of both steps moderate, regardless of whether the belts are split or tied. For example, in the gait on the right side of Fig. 11, the slow belt step length is zero but the swing times for both the fast and slow legs are longer than 0.29 seconds. The extremely short step onto the slow belt does not necessarily require the swing time for either leg to be extremely short.

Robustness and costs associated with lateral balance could also be preventing people from taking full advantage of the treadmill. Although the model gait for people’s chosen fast belt step length is one of the more robust gaits according to our ground height disturbance metric, other aspects of stability might make the model’s gait less desirable than people’s adopted gait. The model’s long fast leg swing time corresponds to a long stance period on the slow belt (Fig. 12A), one that approaches the maximum stance time observed (0.87 sec) when people are asked to walk with the lowest step frequency possible (Nilsson and Thorstensson 1987). To take advantage of the belt speed difference, people would need to significantly extend the amount of time spent on their slow belt leg, and that long single support could increase costs associated with active control of lateral balance (Dean et al. 2007).

Finally, the model gaits with high step length asymmetry would require people to produce a large average free vertical moment during walking (Fig. 13), another possible explanation for why people adopt only a slightly positive step length asymmetry. During normal walking, the average free vertical moment produced by each leg is reportedly between 0.1 (Collins et al. 2009) and 0.9 N · m (Almosnino et al.2009). Altering gait in such a way that the average free vertical moment increases has been associated with increases in energy cost. For example, swinging the arms oppositely from normal results in a higher free vertical moment and a 26% higher metabolic cost of walking (Collins et al. 2009). We estimated the required average free vertical moment for the rimless wheel model by integrating the moment produced about the center of mass by the fore-aft ground reaction forces, adding in the change in angular momentum of the collisions, and dividing the total by the entire cycle time:

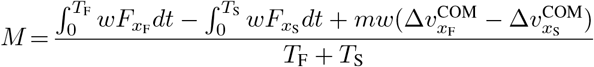

where *T*_F_ and *T*_S_ are the fast and slow leg stance times, *w* is the lateral distance from the contact point to the wheel center (estimated at 6 cm based on human step width of 12% of leg length (Donelan et al. 2001)), *F*_*x*f_ and *F*_*x*S_ are the fore-aft forces during the fast and slow belt rotations respectively, *m* is the total mass, and 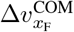 and 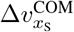 are the change in horizontal velocity of the center of mass during the fast and slow belt collisions respectively. This average moment *M* is created by the ground reaction forces, and people would have to compensate by producing their own free vertical moment in order to avoid turning. Published data do not exist for the average free vertical moment of people’s adapted splitbelt gait. However, at the fast step length that people adopt, our model would require an average free vertical moment higher than that of walking with anti-normal arm swing in Collins et al. (2009) with its associated significant increase in energy cost. Although our model calculation does not include the transverse force contributions to the vertical moment, the high magnitude still suggests that free vertical moment could be an important factor to consider in understanding human split-belt walking. We can imagine that in adapting to the treadmill, people may cease decreasing their slow belt step length once just past symmetry because the increased cost of producing more free vertical moment outweighs the energetic benefit of net work performed by the treadmill.

**Figure 13.**
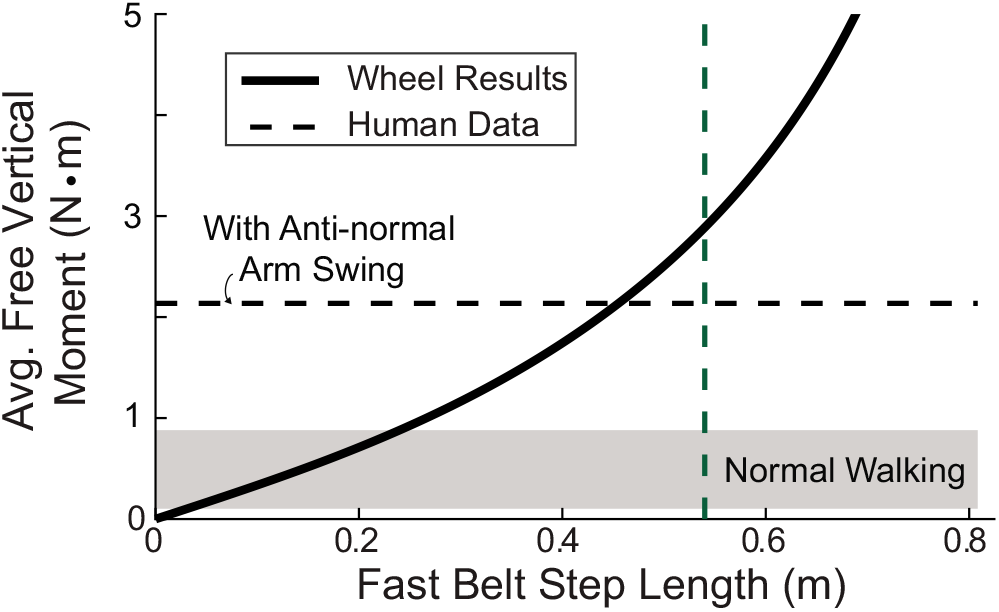
For the family of solutions in Fig. 11, the required average free vertical moment is different for each of the possible gaits. The free vertical moment is one factor which could influence why people choose not to adopt the rimless wheel’s step lengths, since high moments are associated with increased energy costs during walking (Collins et al. 2009). Published data from human experiments for average free vertical moment during walking with normal and anti-normal arm swing are included for comparison (Collins et al. 2009; Almosnino et al.2009).

The comparisons between model gaits and people’s adapted split-belt gait could inform future experiments to further understand what limits people’s ability to take advantage of the split-belt treadmill. While we have not previously observed people being able to lower the cost of split-belt walking below the cost of walking on tied belts at the slow belt speed, it is possible that combinations of belt speeds to facilitate this have simply not yet been tested. We could also enforce different amounts of step length asymmetry for different belt speed combinations to better understand the optimal step length asymmetry and gain insight into how people’s ability to take advantage of the treadmill changes for different speed combinations. To better understand the impact of balance, we could conduct split-belt adaptation experiments in a stabilizing force field, potentially allowing people the freedom to adopt more extreme gaits that better take advantage of the treadmill. We can also imagine attempting to eliminate the effort associated with producing a free vertical moment. Perhaps using a device that allows translation but not rotation would allow people to adopt more asymmetric gaits without having to overcome the tendency to turn. In addition to experiments, adding complexity to the model could prove enlightening for understanding human split-belt treadmill adaptation. Incorporating double support phases, swing costs, pitch and yaw moments, physiologically realistic muscles, and costs associated with negative work performed by the legs could all improve our understanding of the mechanics underlying human split-belt treadmill walking.

The split-belt rimless wheel can also offer insights into rehabilitation techniques for individuals post-stroke. The model is consistent with the practice of using the treadmill to augment step length asymmetry during training in order to reduce asymmetry overground (Reisman et al. 2013). While typically explained as people trying to return to their baseline asymmetry, this lengthening of the fast belt step can also be considered from the model’s energy principle that longer steps onto the fast belt allow people to take advantage of the treadmill. For intervention techniques focused on increasing the stance time of the paretic limb, the model is consistent with the practice of placing the paretic limb on the slow treadmill belt (Malone and Bastian 2014). Rather than considering the intervention as limiting people’s ability to rely on their healthy limb by placing it on the fast belt, this can be viewed as energy optimization driving people toward solutions that rely on longer stance times on the slow belt leg. Finally, the model offers an interesting perspective on using the split-belt treadmill to strengthen and train the paretic limb by forcing people to work harder during rehabilitation. If people were able to take as much advantage of the belt speed difference as suggested by the model, then split-belt walking might actually be easier than tied belt walking, defeating the purposes of the exercise. It may be beneficial for rehabilitation that people are limited in their ability to take advantage of the treadmill to make walking easier.

## 4 A Physical Split-Belt Rimless Wheel

### 4.1 Mechanical Design

We used the simulation results to guide the design of a robot that passively walked forward on a split-belt treadmill (Fig. 14A, B; see also Extension 2, 3). We chose 2*α* = 31°and 2*β* = 9° because those angles demonstrated moderate speed performance and high disturbance rejection in simulation, and their sum divides evenly into 360°. We cut knobs into the edges of two acrylic nonagons to create two identical “rimless wheels” with 9 spokes each (40°spacing between the spokes) and an effective spoke length of 0.254 m. The nonagons were then attached with a 9^°^ angular offset and a 7.6 cm lateral offset, enabling the robot to transition from belt to belt with the desired angles as it alternated steps. We glued small strips of rubber around the ends of the knobs to increase the wheel’s traction on the treadmill during rotations. The mass of the two nonagons and their connecting hardware was 3.53 kg and the radius of gyration was 0.163 m, creating a moment of inertia about the center of mass of 0.094 kg · m^2^.

**Figure 14.**
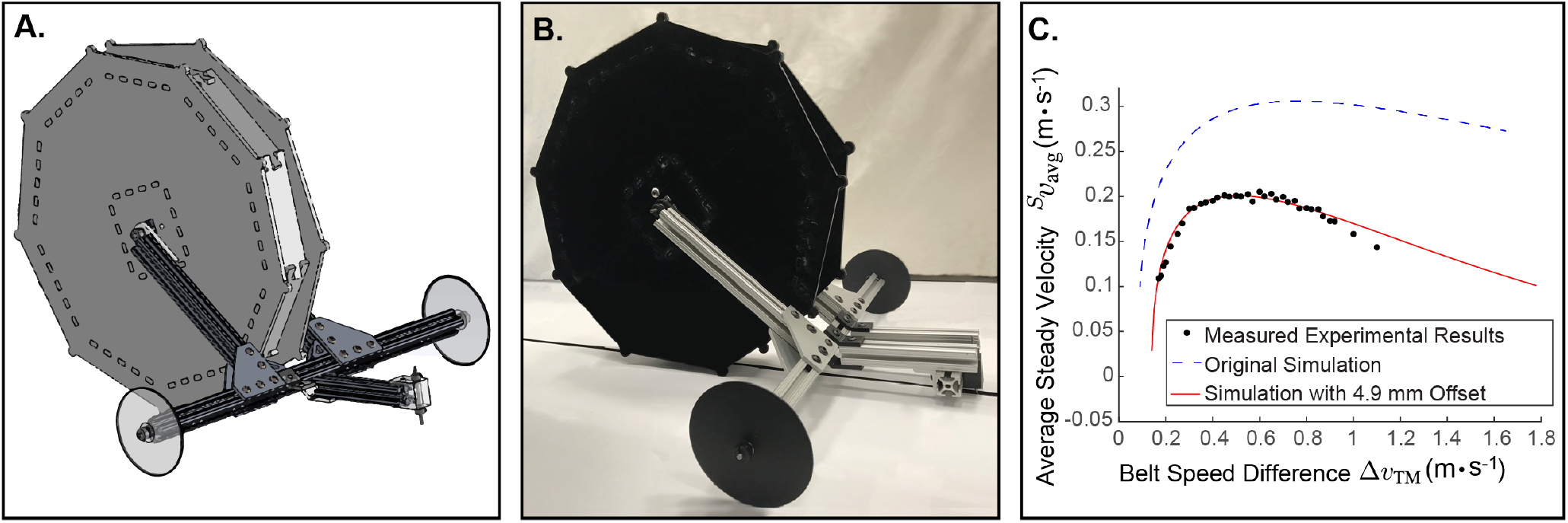
(**A**) CAD model and (**B**) photograph of the split-belt rimless wheel robot (see also Extension 2, 3). (**C**) Results of measuring the split-belt rimless wheel robot’s steady walking speed for a range of treadmill belt speed differences. The simulation results better matched the experimental results when an offset was added between the fast and slow belts in simulation such that the fast belt was 4.9 mm higher than the slow belt.

We added two attachments to eliminate roll and yaw. We placed freely spinning stabilizer wheels at the front to prevent the robot from leaning side-to-side during rotation and prematurely contacting the opposite belt. The stabilizer wheels were positioned such that the robot’s center of mass always remained within the triangle of support created by the two wheels and whichever knob was in contact with the ground at that instant. A small rudder, placed in front of the stabilizer wheels, rolled in bearings along the gap between the belts to counteract the vertical moment and prevent the robot from turning as it walked forward.

### 4.2 Experimental Comparisons

To test the robot’s ability to walk on a split-belt treadmill and to validate the simulation, we measured the robot’s slow-belt walking speed for a range of belt speed differences (Fig. 14C). We held the slow belt stationary and set the fast belt to speeds ranging from 0.15 to 1.1 m · s^−1^. For each belt speed difference, we allowed the wheel several steps to stabilize and then timed it walking forward a pre-determined distance. We estimated the speed in the slow belt reference frame as the pre-determined distance divided by the time. We completed at least five trials per belt speed difference and averaged the trials to obtain a final velocity.

We compared the experimental results to results from a simulation model adjusted for size, mass, rotational inertia, and collision angles of the physical robot (Fig. 14C). The results were qualitatively similar, with both the physical robot and the simulation demonstrating a rapid increase and then slower decrease in steady walking speed as the belt speed difference increased. However, the physical prototype consistently walked more slowly than the simulation model.

We considered various adjustments to the simulation model to determine which differences between the real world and the simulation could be responsible for the offset in walking speed. Friction and general inefficiencies in the system, modelled as an incline, did not explain the difference, nor did variations in leg length, inertia, or collision angles. Random variations in the ground height slowed the simulated wheel but not in a way consistent with the experimental results. The modification that best brought the simulation results into agreement with the experimental results was setting the fast treadmill belt 4.9 mm higher than the slow treadmill belt (Fig. 14C).

Although the belts were not physically offset, the physical wheel may have been interacting differently with the two belts, and the simulated 4.9 mm offset may have captured some of the asymmetries observed experimentally. For example, the robot appeared to bounce upon contact with the fast belt but not with the slow belt. The rubber at the edges of the robot could also have interacted differently with the two belts, taking a moment longer at fast belt contact to grip the belt firmly and begin rotation. With a more complex contact model, or other complicating features, the simulation model could have been brought into better agreement with experimental results. However, despite the simplified physics used, the simulation model made predictions that were qualitatively consistent with the real system. Most importantly, the physical robot was successful in demonstrating that a split-belt rimless wheel can harness energy from a treadmill to roll steadily on level ground.

### 4.3 Comparisons to Traditional Energy Capture from Dual-Velocity Environments

The rimless wheel takes advantage of external energy differently than traditional machines like sailboats, water wheels, and land yachts, demonstrating a new way that devices can harness energy from relative motion in the environment. Traditional mechanisms rely on continuous contact with both substrates of different velocity, but this wheel alternates contact between the fast and slow belts. The wheel also requires a process of energy capture, exchange, and storage not present in traditional systems. The wheel harvests energy during the fast belt rotation by converting treadmill energy into gravitational potential energy and then turning it back into kinetic energy during the slow belt rotation. The split-belt wheel’s angular velocity must change throughout the gait cycle as energy is captured and converted, unlike traditional devices that can operate at constant speeds with no need to store or exchange energy.

The split-belt rimless wheel’s method of energy harnessing is mechanically much more similar to that of an albatross during dynamic soaring. Both the wheel and the bird spend some time in a faster reference frame where they convert kinetic energy to potential energy at an advantageous rate, and the longer they spend in the faster frame, the more energy they capture. One key difference is the asymmetry enabling that energy capture: the wheel relies on unequal step lengths onto the fast and slow belts whereas the bird relies on a change in its heading angle during ascent and descent. Additionally, the wheel abruptly alternates between the two treadmill belts, with losses due to collisions, whereas the albatross moves smoothly between the wind layers, with losses due to drag. It is worth noting, however, that the albatross model of energy capture would still work for an instant change in airspeed.

This mechanism for energy capture from dual-velocity environments could be exploited in other contexts. Existing dual-velocity devices could be modified to incorporate asymmetric, intermittent contacts to allow greater energy capture. New applications could be discovered, for example in robots with aerospace or transportation applications. We can imagine, for instance, employing this concept to move materials in a factory in the direction opposite the motion of a conveyor belt. This work also suggests a broader class of energy harvesting mechanisms, applicable to any robot that does not operate within a single Newtonian reference frame.

## 5 Conclusions

In this study, we found that a split-belt rimless wheel, simulated or physical, can harness enough energy from treadmill belts moving at different speeds to walk steadily under a variety of conditions. When the collision angles are appropriately asymmetric to take advantage of the belt speed difference, higher fast belt speed can actually be beneficial; making the fast belt move backwards faster increases the wheel’s forward speed and improves its disturbance rejection.

Our simulation work helps build a framework to better understand the mechanics underlying human split-belt treadmill walking. Without an impact of muscles or training, our model demonstrates how split-belt walking may require less mechanical work than tied-belt walking at the slow belt speed. The asymmetry in step lengths enabling the treadmill to perform work on the wheel offers insight into why people adopt a positive step length asymmetry after a period of split-belt adaptation. More complicated simulation models, as well as future human-subject experiments, can further reveal the opportunities and limitations for humans to take advantage of the split-belt treadmill.

Our physical prototype demonstrates the robustness of the split-belt rimless wheel’s energy capture mechanism, illustrating possibilities for transfer to other devices. The wheel can harness energy under a range of circumstances and withstand speed and ground height disturbances, despite imperfections in construction. Future work with a physical split-belt rimless wheel could include improving the model with better contact modeling or creating a modular prototype to enable expanded physical validation of the simulation model, further clarifying how this mechanism might best be implemented in other devices.

## Supporting information

Wheel Simulations

Physical Wheel Videos

## Acknowledgements

The authors thank Rachel Adenekan and Katherine Poggensee for assistance with experimental data collection.

## 6 Appendices

### 6.1 Appendix A: Index to Multimedia Extensions

| Extension | Media Type | Description |
| --- | --- | --- |
| 1 | Video | Simulated wheel walking on split-belt treadmill |
| 2 | Video | Physical split-belt rimless wheel on tied belts and split belts |
| 3 | CAD | CAD assembly and parts for the physical split-belt rimless wheel robot. |

### 6.2 Appendix B: Derivation of System of Equations to Solve for Steady Walking

In the main text, we explain how each collision and rotation of the gait cycle has a governing equation relating the angular velocity before the event to the angular velocity after it. These four governing equations become a system of equations that can be solved for the fixed point angular velocity at the beginning of the slow belt rotation, 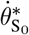. In this section, we show the complete derivation of those four equations. As in the rest of the text, the subscript S_0_ refers to the beginning of the slow belt rotation, S_f_ the end of the slow belt rotation, F_0_ the beginning of the fast belt rotation, and F_f_the end of the fast belt rotation. The left-superscripts *S* and *F* indicate whether a frame-dependent term is expressed in the fast or slow belt reference frame.

#### 6.2.1 Rotation on the Slow Treadmill Belt

Given the velocity at the beginning of the slow belt rotation, the velocity at the can be determined using an energy balance approach (Eq. 1).

Before the rotation, the wheel’s velocity in the slow belt reference frame is

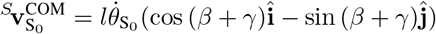

where 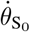 is the angular velocity before the rotation. The total energy is a combination of the kinetic and potential energies,

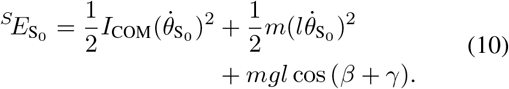

After the rotation, the wheel’s center of mass velocity in the slow belt reference frame is

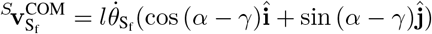

where 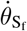 is the angular velocity after the rotation. The total energy after rotation is a combination of the kinetic and potential energies,

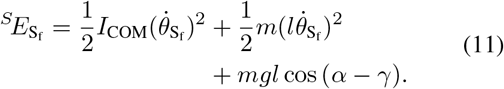

Viewed in the slow belt reference frame, the wheel behaves like an inverted pendulum rotating over a stationary contact point. It exchanges potential and kinetic energy during the rotation, but the total energy remains constant. We set the energies 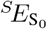 and 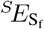 (Eqs.10 and 11) equal,

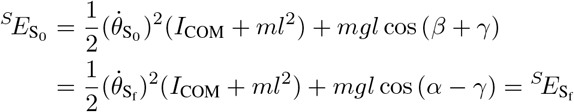

and rearrange to solve for the angular velocity after rotation 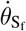 as a function of the angular velocity before rotation 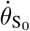,

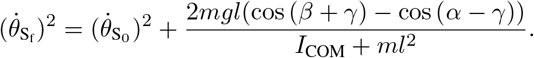

#### 6.2.2 Collision onto the Fast Treadmill Belt

Conservation of angular momentum about the new foot contact, here computed in the slow belt frame, can be used to determine the angular velocity after the fast belt collision 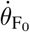 (Eq.4).

The angular momentum **H** in general terms is

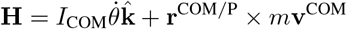

where **r**^COM/P^ is the position vector from the point colliding with the fast treadmill belt to the center of mass and **v**^COM^ is the center of mass velocity. The position vector is

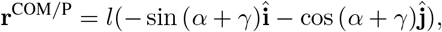

and it is the same both before and after the fast belt collision.

The angular momentum before the collision is

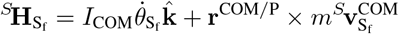

where the center of mass velocity expressed in the slow belt reference frame is

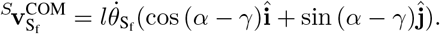

The angular momentum before the collision, 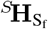, when expressed in the slow belt reference frame, simplifies to

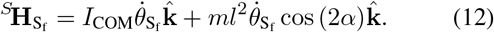

The angular momentum after the collision is

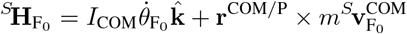

where the center of mass velocity expressed in the slow belt reference frame is

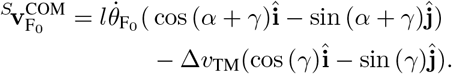

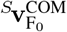 includes the belt speed difference Δ*ν*_TM_ when expressed in the slow belt frame because the wheel at that point is rotating on a fast belt spoke. In the fast belt frame, the center of mass speed would just be 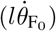. Because the angular momentum after the collision must be computed in the slow belt frame to be equated with the angular momentum before the collision, 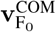 must be in the slow belt frame when used.

The angular momentum after the collision 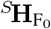, when expressed in the slow belt reference frame, simplifies to

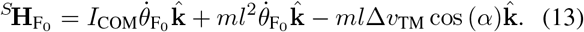

We set the angular momenta 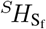 and 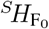 (Eqs.12 and 13) equal through the conservation of angular momentum,

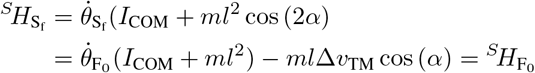

and rearrange to solve for the angular velocity after the collision 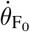 as a function of the angular velocity before the collision 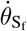,

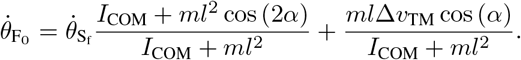

#### 6.2.3 Rotation on the Fast Treadmill Belt

The fast belt rotation is similar to the slow belt rotation, and an energy balance is used to find the angular velocity post-rotation, but the energy balance is computed in the fast belt reference frame (Eq. 5). Viewed in the fast belt reference frame, the contact point between the wheel and the treadmill appears stationary during the rotation, making the wheel an inverted pendulum rotating over a stationary contact point and maintaining constant energy.

Before the rotation, the wheel’s velocity in the fast belt reference frame is

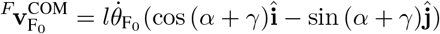

where 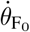 is the angular velocity before the rotation. The total energy is a combination of the kinetic and potential energies,

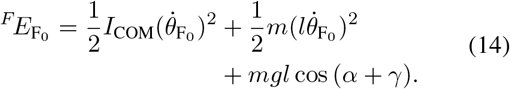

After the rotation, the wheel’s velocity in the fast belt eference frame is

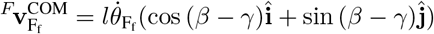

where 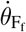 is the angular velocity after the rotation. The total energy is a combination of the kinetic and potential energies,

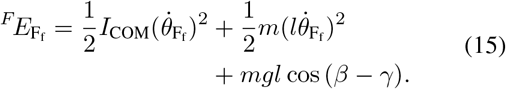

We set the energies 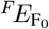 and 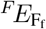 (Eqs.14 and 15) equal,

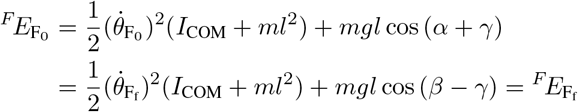

and rearrange to solve for the angular velocity after rotation 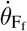 as a function of the angular velocity before rotation 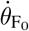,

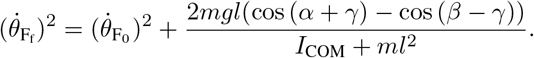

#### 6.2.4 Collision onto the Slow Treadmill Belt

The slow belt collision is similar to the fast belt collision, with a conservation of angular momentum computed in the slow frame being used to relate the angular velocity before and after the collision (Eq. 8).

The angular momentum **H** in general terms is still

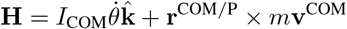

but the position vector **r**^COM/P^ is now

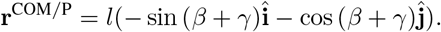

The angular momentum before the collision is

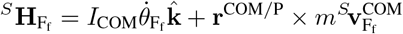

where the center of mass velocity expressed in the slow belt reference frame is

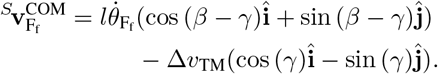

The velocity includes the belt speed difference Δ*ν*_TM_ because the wheel rotates on a fast belt spoke before the collision, but the velocity must be expressed in the slow belt reference frame to compute the angular momentum in the slow belt frame.

The angular momentum before the collision, 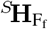, when expressed in the slow belt reference frame, simplifies to

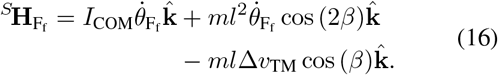

The angular momentum after the collision is

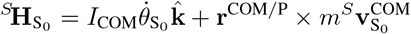

where the center of mass velocity expressed in the slow belt reference frame is

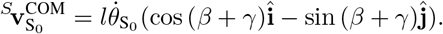

The angular momentum after the collision 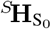, when expressed in the slow belt reference frame, simplifies to

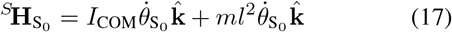

We set the angular momenta 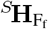 and 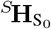 (Eqs.16 and 17) equal through the conservation of angular momentum,

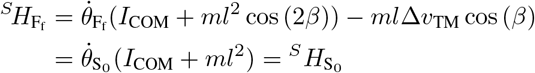

and rearrange to solve for the angular velocity after the collision 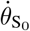 as a function of the angular velocity before the collision 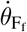,

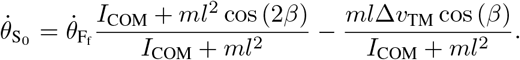

#### 6.2.5 Solving the System of Equations

Together, the four governing equations for a gait cycle are

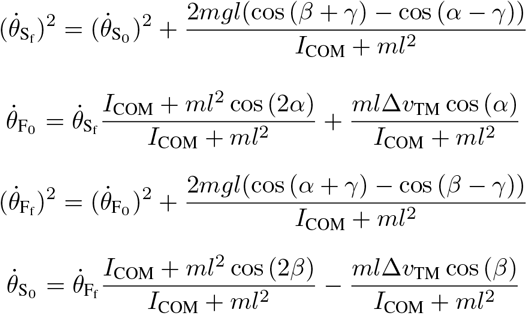

where the four unknowns are 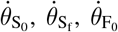, and 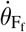. These can be combined into one equation containing any one of the variables. We combined the equations into 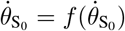 and solved to find the fixed point angular velocity 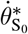.

There are numerical solutions where the angular velocity is negative at some point in the cycle, indicating that the wheel is falling backward, but the equation solves because the angular velocity is squared when appearing in the equation. We enforced that the angular velocity be positive in all cases.

### 6.3 Appendix C: Fast Belt Rotation Potential Energy for Non-zero Treadmill Incline

When the treadmill incline *γ* is non-zero, the wheel’s potential energy during the fast belt rotation changes as the fast belt moves down relative to the slow belt during the rotation (Fig. 15). We compute the potential energy relative to the height of the contact point between the wheel and the treadmill during the slow belt rotation, even though the wheel rotates on a different, moving contact point during the fast belt rotation.

During the slow belt rotation, the potential energy in the slow belt reference frame is

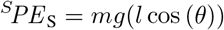

where *θ* is the angle from vertical 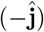 to the contacting spoke. At the end of the slow belt rotation, the wheel collides with the fast treadmill and the contact point is redefined as the end of the fast belt spoke. Redefining the contact point changes the value of *θ*, so the quantity (*l* cos(*θ*)) changes. However, the wheel’s center of mass does not change position in the instant of the collision, so the potential energy should not change.

**Figure 15.**
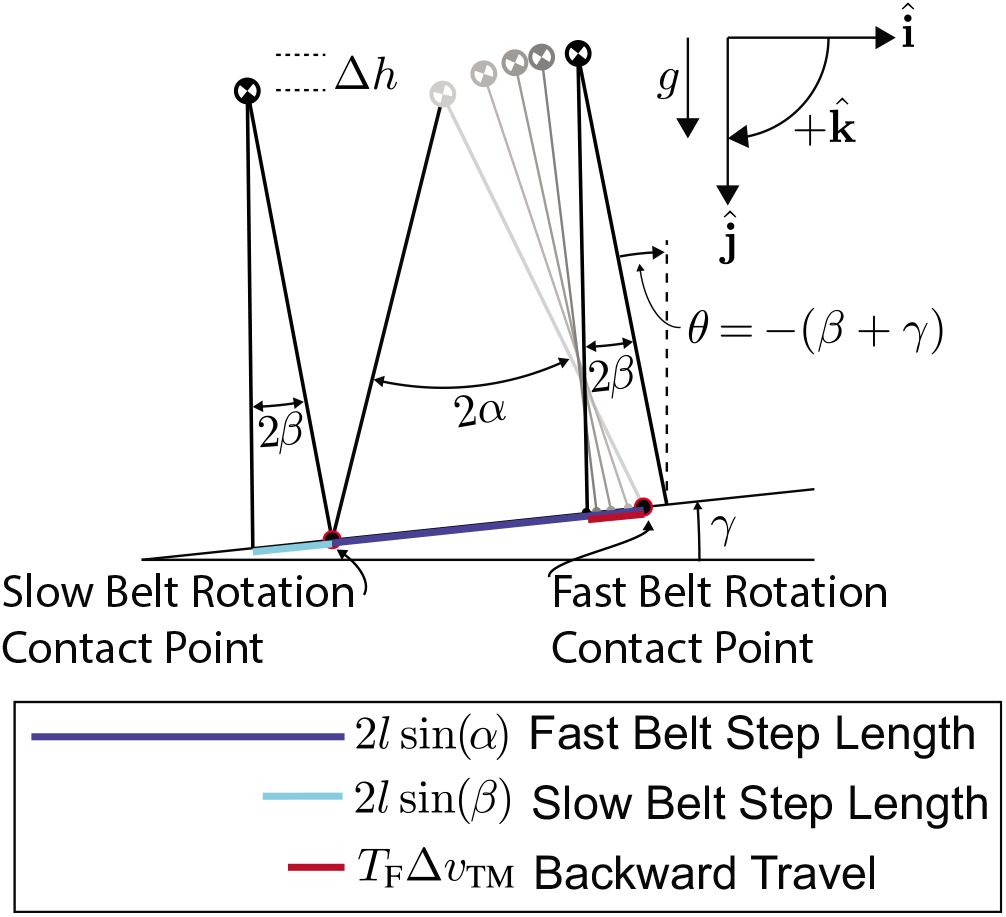
When the wheel walks up an incline *γ*, the amount the center of mass moves up during a gait cycle Δ*h* is a function of the fast belt step length 2*l* sin (*α*), the slow belt step length 2*l* sin (*β*), and the amount the wheel moves backward during the fast belt rotation *T*_f_Δ*ν*_TM_.

When calculating the potential energy during the fast belt rotation, we add a term to compensate for the difference in height of the contact point before and after the collision. Just after the collision onto the fast belt, the wheel’s potential energy in the slow belt frame is

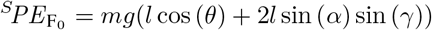

where 2*l* sin (*α*) sin (*γ*) is the difference in height of the two contact points during the collision.

As the wheel rotates on the fast treadmill belt, the fast belt moves down relative to the slow belt. To track the potential energy relative to the original slow belt contact point, we must account for the downward motion of the wheel. We add another term to the expression for the potential energy so that potential energy during the fast belt rotation is

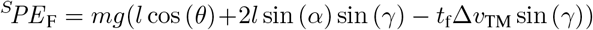

where *t*_F_ is the time since the fast belt collision. At the end of the fast belt rotation, the wheel collides with the slow treadmill belt and the contact point is again redefined. We would need to account for that redefinition to continue tracking the wheel’s potential energy into another gait cycle, but we complete all analysis in just one representative cycle. Subtracting the wheel’s potential energy at the beginning of the slow belt rotation 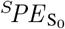 from the final potential energy at end of the fast belt rotation 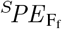 gives the change in potential energy over one gait cycle 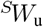.

For uphill walking, 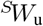 is a crucial result that defines whether the wheel can walk steadily. Substituting in for the angles at the beginning of the slow belt rotation and the end of the fast belt rotation gives us a specific equation for the energy change over a cycle. At the beginning of the slow belt rotation, *θ* = −(*β* + *γ*) and

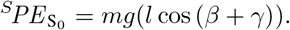

At the end of the fast belt rotation, *θ* = (*β* −*γ*) and

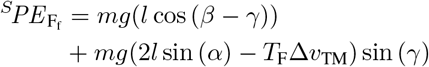

where *T*_F_ is the duration of the fast belt rotation. The net energy change during one uphill gait cycle is

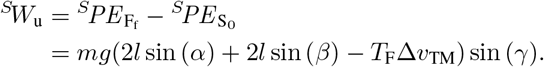

The 2*l* sin (*α*) sin (γ) and 2*l* sin (*β*) sin (γ) terms account for how much the center of mass would rise if the belts were stationary, and the −*T*_F_Δ*ν*_TM_sin (γ) term accounts for the downward motion of the fast belt relative to the slow belt.

### 6.4 Appendix D: Impact of Changing Belt Speed Difference, Fast Belt Collision Angle, and Slow Belt Collision Angle on Collision Losses and Rotation Gains

The collision angles and belt speed difference affect the energy lost in collisions and gained in the fast belt rotation. There is nuance because the components of the cycle are so interconnected, but we can still get a general picture of the energy effect by looking at the collisions and rotations individually as we change different parameters.

Increasing the belt speed difference decreases the energy lost in the fast belt collision 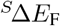 (Fig. 16A) but increases the energy lost in the slow belt collision 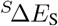 (Fig. 16B). Because the angle between the spokes is larger in the fast belt collision, the decrease in ^S^ΔEF is larger than the increase in 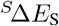 so that increasing the belt speed difference is overall beneficial from an energy standpoint.

**Figure 16.**
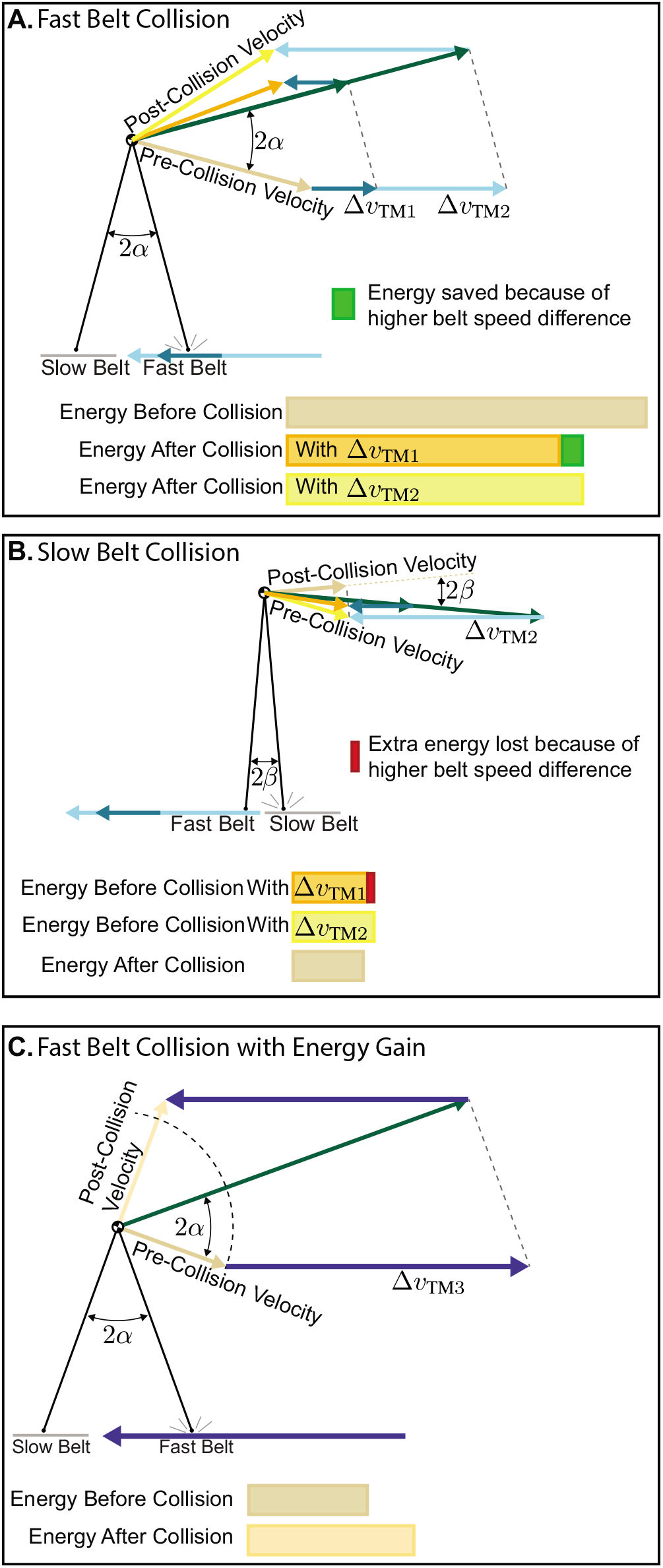
**A**. Increasing the belt speed difference Δ*ν*_TM_ decreases the energy lost in the fast belt collision. **B**. Increasing Δ*ν*_TM_ increases the energy lost in the slow belt collision. The wheel would be walking steadily for these angular velocities with Δ*ν*_TM1_. By increasing from Δ*ν*_TM1_ to Δ*ν*_TM2_, the savings in the fast belt collision (green box) are larger than the added costs in the slow belt collision (red box), so increasing Δ*ν*_TM_ is overall beneficial. **C**. There are cases where Δ*ν*_TM_ is high enough that the wheel actually gains energy in the fast belt collision when viewed in the slow belt reference frame.

It is possible to increase the belt speed difference so much that energy is actually gained in the fast belt collision (Fig. 16C). This never happens in the simplest model variation where the slow belt collision angle is zero but can happen when *β* is non-zero. The energy change for the cycle must be zero when the wheel walks on level ground, and energy is always gained during the fast belt rotation. In the case where *β* is zero, there is no energy change in the slow belt rotation or collision, so energy must be lost in the fast belt collision. When *β* is non-zero, some energy is lost in the slow belt collision, so energy can be gained in both the fast belt rotation and the fast belt collision with the total system energy still equaling zero over the cycle as long as enough energy is lost in the slow belt collision.

Increasing the belt speed difference also increases the energy gained during the fast belt rotation 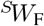 when the angular velocity at the start of the fast belt rotation is the same. This can be explained with the same work equation used to explain how the treadmill can perform any work. The belt speed difference increases the contact point velocity 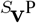, while the force **F**, dependent on the wheel’s angular position and velocity, remains unchanged. When the wheel rotates up with the same angular velocity, the work for the rotation, 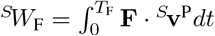 is higher because 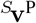 increases while **F** and *T*_F_ remain the same.

Decreasing the collision angles decreases the energy lost in the collisions. This is true regardless of the belt speed difference. When the angles are larger, the center of mass velocity must be more dramatically redirected, meaning more energy is lost. As the angle decreases, less energy is lost until no energy is lost at the extreme when the angle between the spokes at the collision is infinitesimally small.

The impact of changing the collision angles on the work gained in the fast belt rotation 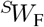 is different for the two angles. Increasing *α* at the fast belt collision increases 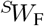 while increasing *β* decreases 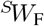. The power is positive as the wheel rotates up through *α* from the fast belt collision to vertical. When *α* is larger, the wheel can extract more energy from the treadmill. Once the wheel passes vertical, the power is negative as the wheel rotates down through *β*. The larger *β* is, the longer the wheel spends rotating down with the treadmill doing negative work and removing energy.

Whether changing a parameter causes the wheel to rotate faster or slower depends on how that parameter impacts both collision losses and rotation gain. Increasing the belt speed difference Δ*ν*_TM_ allows the wheel to rotate faster because the wheel loses less energy in the collisions and gains more energy in the rotation. Increasing the slow belt collision angle 2*β* forces the wheel to rotate more slowly because the wheel loses more energy in the slow belt collision and gains less energy in the fast belt rotation. When changing the fast belt collision angle 2*α*, the overall result is less obvious. Decreasing *α* decreases the fast belt collision loss but also decreases the energy gained in rotation. It turns out that the decrease in collision loss is larger than the decrease in rotation gain because the decrease in rotation gain is nearly linear while the decrease in collision loss is close to quadratic. The result is a net benefit such that the wheel can rotate more quickly for a smaller *α*. As *α* continues to decrease, the gap between the change in collision loss and the change in rotation gain decreases such that decreasing *α* has a smaller impact when *α*is already small than when it is large.

### 6.5 Appendix E: Increases in Angular Velocity Often Increase Slow-Belt Walking Speed

We showed that changing the collision angles and belt speed difference changes the angular velocity with which the wheel must walk. If the parameters changed but the wheel walked with the same angular velocity, it would speed up or slow down because the energy change over the course of a cycle would be non-zero. The wheel’s angular velocity and its slow-belt walking speed 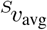 are closely related but changing a parameter in a way that causes an increase in angular velocity does not always increase the wheel’s walking speed.

Recall that the equation for the wheel’s steady walking speed 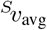 is

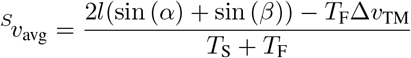

where 2*l* sin (*α*) is the last belt step length, 2*l* sin (*β*) is the slow belt step length, *T*_S_ and *T*_F_ are the durations of the slow and fast belt rotations, and T_F_Δ*ν*_TM_ is the backward distance the wheel travels while riding on the fast belt.

Increasing the belt speed difference Δ*ν*_TM_ always increases the wheel’s angular velocity because the wheel loses less energy in collisions and gains more energy in the fast belt rotation. That increase in angular velocity decreases both *T*_S_ and *T*_F_, which should increase 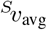. However, the belt speed difference increase also means the wheel travels backward faster during the fast belt rotation. Even though *T*_F_ decreases when Δ*ν*_TM_ increases, the backward distance *T*_F_Δ*ν*_TM_ tends to increase as Δ*ν*_TM_ increases. Since both the numerator and denominator decrease, the wheel’s slow-belt walking speed can increase or decrease depending on which aspect dominates. Most of the time, the decrease in rotation durations outweighs the increased backward distance so that increasing Δ*ν*_TM_ increases 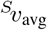. However, this is not guaranteed, and sometimes the increase in Δ*ν*_TM_ actually decreases 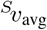 despite increasing the wheel’s angular velocity.

Except when the wheel walks uphill, decreasing the fast belt collision angle always increases the wheel’s angular velocity. The wheel rotates faster and rotates through a smaller angle, decreasing both *T*_S_ and *T*_F_, and decreasing *T*_F_Δ*ν*_TM_. However, the fast belt step length 2*l* sin(*α*) is also smaller when *α* decreases. The lower rotation durations and smaller backward distance during the fast belt rotation outweigh the shorter step length, and the angular velocity increase associated with decreasing *α* always increases 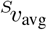.

Decreasing the slow belt collision angle 2*β* also increases the wheel’s angular velocity and increases 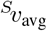. Like with the fast belt collision angle, decreasing *β* allows the wheel to walk with higher angular velocity, decreasing both rotation durations *T*_S_ and *T*_F_ and decreasing the backward distance *T*_F_Δ*ν*_TM_. The slow belt step length 2*l* sin(*β*) decreases, but the other terms dominate and 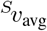 increases when *β* decreases.

### 6.6 Appendix F: Eigenvalue Stability Analysis

The eigenvalue for the discrete, linearized stride-to-stride state transition matrix is always positive and less than or equal to one. An eigenvalue of one indicates that all of the angular velocity perturbation is still present after one gait cycle, and the wheel rolls steadily with the new angular velocity. This only occurs when both collision angles are infinitesimally small and the rimless wheel is essentially a rimmed wheel (Fig. 17). A theoretical rimmed wheel can roll steadily with any angular velocity, so this is a logical result.

**Figure 17.**
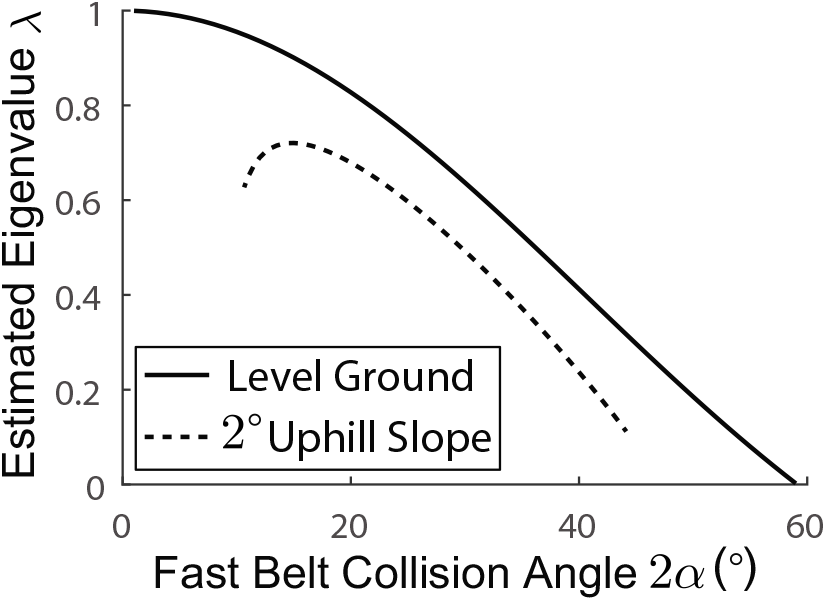
Eigenvalue stability analysis for a few representative cases. The results shown here are for a slow belt collision angle and moment of inertia about the center of mass of zero. The treadmill belt speed difference was 2.0 m · s^−1^ for the level ground case shown and 1.0 m · s^−1^ for the uphill case. According to this metric, the wheel is more stable when the eigenvalue is smaller and the wheel more quickly rejects disturbances. An eigenvalue of one indicates that the wheel does not dissipate any of the perturbation, while an eigenvalue of zero indicates that the wheel returns to its steady walking speed within just one gait cycle.

At the other extreme, an eigenvalue of zero indicates that the entire perturbation disappears and the wheel returns to the fixed point angular velocity by the end of one gait cycle. The wheel approaches this result in the extreme case where the slow belt collision angle 2*β* is infinitesimally small, and the fast belt collision angle 2*α* is at its maximum feasible value for a given belt speed difference. In this case, the wheel’s angular velocity after the collision onto the slow belt is approximately zero, indicating that the wheel must roll as slowly as possible to gain enough energy from the fast belt during rotation. A perturbation which increases the wheel’s angular velocity cannot be sustained and the wheel returns to the fixed point very quickly. For these extremely wide angles, the angular velocity perturbation must be positive because a negative perturbation can cause the wheel to fall backward, failing to complete the cycle.

According to this metric, the split-belt rimless wheel is generally more stable with larger angles. An angular velocity perturbation is more quickly dissipated and the wheel approaches its steady-state walking velocity in a smaller number of steps when it has larger collision angles. The notable exception to this behavior is when there is an uphill slope. When walking uphill, a wheel with the smallest possible feasible collision angles seems more stable than one with slightly larger collision angles. As the collision angles continue to increase, however, the normal pattern returns and increasing the angles increases the stability (Fig. 17).

### 6.7 Appendix G: Ground Height Disturbances with Small Collision Angles

During the variable terrain simulations, we found that a wheel with small collision angles often failed to complete rotations and fell backward. This is because a step of the same height has a larger impact on a wheel with small collision angles (Fig. 18). The fast belt collision angle 2*α*defines how much the wheel rotates up during the fast belt rotation and how much energy is gained in the rotation. When the wheel takes a step down onto the fast belt, it rotates less on the fast treadmill belt and gains less energy. When *α* is the larger *α*_1_ the lost upward rotation Δ_1_ is smaller than the lost rotation Δ_2_ with *α*_2_. With *α*_2_, the wheel barely rotates up at all during the fast belt rotation and ends up losing energy during the rotation because the negative work from rotating down through *β* is larger than the positive work from rotating up. This phenomenon helps illustrate why a wheel with small fast belt collision angles is not as tolerant to ground height disturbances as one with intermediate collision angles.

**Figure 18.**
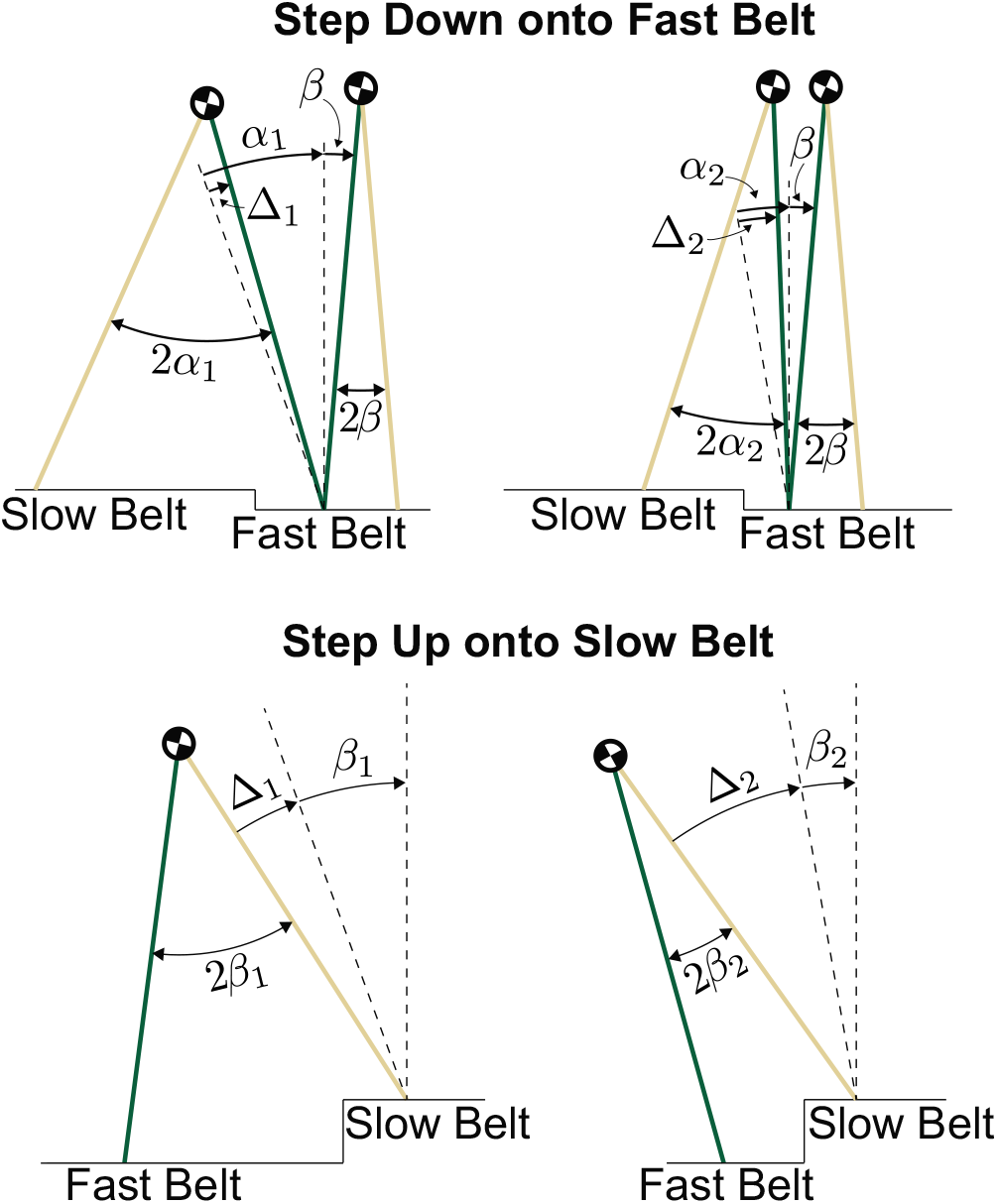
Small collision angles make the wheel less tolerant to ground height disturbances because a step of the same size has a larger impact on a wheel with small angles. When the wheel steps down onto the fast treadmill belt, the amount of energy captured from the treadmill decreases because the wheel rotates up through a smaller angle (*α* −Δ) and Δ is larger when the angle is smaller. When the wheel steps up onto the slow treadmill belt, it must rotate up through (*β*+Δ) to reach vertical. For small collision angles, the added amount of rotation is large and can cause the wheel to fall backward.

Steps up onto the slow treadmill belt demonstrate the large impact of a step when the slow belt collision angle 2*β* is small. In order to continue walking after a step up, the wheel must be able to reach the top of every rotation. When the wheel takes a step up onto the slow belt, the amount it must rotate up to reach vertical increases. When βis smaller, the extra rotation due to a step up is larger (Δ_2_ > Δ_1_). Even if the wheel had a sizeable margin between its typical angular velocity after a step onto the slow belt and the minimum angular velocity required to reach the top, a step up causing a large Δcould cause the wheel to fall backward.

The step up onto the slow belt also affects the amount of time the wheel spends on the fast belt. When the wheel takes a step up that is small compared to the step length (*β*_1_), the wheel actually gets more work from the treadmill during the fast belt rotation because it does not rotate as far past vertical before the slow belt collision. On the other hand, when the step up is large compared to the step length (*β*_2_), the wheel’s angle at the collision changes more dramatically. The wheel no longer rotates all the way up to vertical during the fast belt rotation, instead colliding with the slow belt very early into the fast belt rotation, limiting the amount of work the wheel can capture from the treadmill. Small slow belt collision angles make the wheel less tolerant to ground height disturbances because the wheel is highly susceptible to falling backward.

